# Soluble guanylate cyclase stimulation mitigates skeletal and cardiac muscle dysfunction in a mdx model of Duchenne muscular dystrophy

**DOI:** 10.1101/2021.02.14.431156

**Authors:** Ling Zhang, Yuanyuan Xu, Keyvan Yousefi, Camila I. Irion, Roger A. Alvarez, Shalini M. Krishnan, Johannes-Peter Stasch, Eliana C. Martinez, Emmanuel S. Buys, Peter Sandner, Lina A. Shehadeh, Justin M. Percival

## Abstract

The impairment of neuronal nitric oxide synthase (nNOS) signaling contributes to disease pathology in the muscle wasting disorder Duchenne muscular dystrophy (DMD). nNOS signal propagation occurs through nitric oxide sensitive soluble guanylate cyclase (sGC), a critical source of cyclic guanosine monophosphate (cGMP) in muscle. Although both nNOS and sGC activity are impaired in DMD patients, little is known about sGC as a therapeutic target. In this study, we tested the hypothesis that stimulating sGC activity with the allosteric agonist BAY41-8543 mitigates striated muscle pathology in the mdx4cv mouse model of DMD. In contrast to DMD patients, mdx mice exhibited greater basal sGC activity than wild type controls with preservation of cGMP levels due partly to upregulation of sGC in some muscles. Stimulating sGC activity in mdx mice with BAY41-8543 substantially reduced skeletal muscle damage, macrophage densities and inflammation and significantly increased resistance to contraction-induced fatigue. BAY41-8543 also enhanced *in vivo* diaphragm function while reducing breathing irregularities suggesting improved respiratory function. BAY41-8543 attenuated cardiac hypertrophic remodeling, fibrosis and diastolic dysfunction including left atrium enlargement in aged mdx mice. Overall, sGC stimulation significantly mitigated skeletal and cardio-respiratory dysfunction in mdx4cv mice. Importantly, this study provides compelling pre-clinical evidence supporting sGC as a novel target in DMD and the repurposing of FDA-approved sGC stimulators, such as riociguat and veraciguat, as a novel therapeutic approach for DMD.

## Introduction

Duchenne muscular dystrophy (DMD) is an X-linked genetic disorder caused by dystrophin gene mutations with a prevalence of 1 in every 7,250 males aged 5 to 24 years [1-3]. These mutations prevent the expression of dystrophin protein causing progressive muscle wasting and weakness, which culminates in the loss of walking ability and ultimately respiratory failure in most cases. The loss of dystrophin can also cause heart failure, an increasing cause of mortality in DMD due to improvements in supportive care. Cardiomyopathy is also a clinical concern in Becker muscular dystrophy (BMD) caused by more benign mutations in the dystrophin gene [4]. At present, there is no cure for DMD. Advances in palliative care, including the use of improved ventilatory support and glucocorticoids, have improved disease course and survival in DMD. However, steroids are effective for a limited time and have considerable side effects. Exon skipping drugs are an approved therapy that may provide benefit in some patients [5-8]. Thus, an urgent need remains for therapies that mitigate pathology in all DMD and BMD patients.

One promising therapeutic strategy for both DMD and BMD is to restore the pathogenic inhibition of neuronal nitric oxide synthase (nNOS) signaling [9, 10]. In the body, nNOS synthesizes the majority of nitric oxide (NO) and plays key roles in neurons and striated muscle cells that are critical to viability in mice and humans [11-15]. The loss of dystrophin decreases expression of nNOSμ and prevents α-syntrophin-mediated recruitment of nNOSμ to the sarcolemmal dystrophin protein complex [10, 16-20]. The loss of dystrophin may also impair the localization of the nNOSβ splice variant further disrupting NO signaling in striated muscles [21, 22]. The loss of nNOS activity exacerbates dystrophic pathology by causing blood delivery defects, skeletal muscle weakness, inflammation and fatigue [18, 21, 23-25]. Accordingly, genetic approaches that increase nNOS activity or restore sarcolemmal nNOSμ in mdx mouse models of DMD restore blood delivery in skeletal muscles and alleviate striated muscle damage and weakness [25-28].

The importance of restoring nNOS in dystrophic muscle is further supported by preclinical gene therapy studies and is a major difference between candidate microdystrophin-based gene therapies being tested clinically [29]. Microdystrophin cassettes that restore sarcolemmal nNOSμ provide greater therapeutic benefit to skeletal and cardiac muscle in mdx mice than microdystrophins that do not [18, 30]. It is important to note that interventions that restore nNOS signaling could be effective as a palliative stand-alone therapy or be used in combination with exon skipping and microdystrophin-based approaches that cannot restore nNOS signaling to enhance their therapeutic efficacy.

nNOS exerts many of its biological functions through its “receptor” soluble guanylate cyclase [31]. nNOS synthesizes NO that binds and stimulates sGC to synthesize cGMP. In turn, cGMP activates protein kinase G, ion channels and phosphodiesterases that mediate NO-cGMP signal transduction. nNOS is required for sGC activity and regulation in skeletal muscle [32]. Accordingly, due to the loss of nNOSμ, individuals with DMD and BMD have substantial reductions in skeletal muscle cGMP synthesis indicating impaired sGC activity [33]. Inhibition of sGC causes skeletal muscle fatigue and increases the risk of myocardial infarction in humans [32, 34]. Therefore, a substantial body of evidence supports a key role for nNOS-sGC-cGMP signaling in cardiac and skeletal muscle dysfunction in DMD and BMD.

The emergence of potent small molecule agonists for sGC has provided a powerful new approach to restore nNOS-sGC-cGMP signaling in DMD and BMD [35]. Agonists such as riociguat, vericiguat, olinciguat and praliciguat are sGC stimulators that bind to native heme-containing sGC and promote cGMP synthesis [36-39]. sGC stimulators can work independently of NO and in synergy with endogenous NO by allosteric stabilization of NO-sGC binding. Therefore, sGC stimulators are less dependent on NO and cGMP levels and thus avoid a key limitation of phosphodiesterase 5 (PDE5) inhibitors that have had limited success in DMD and BMD patients [40-42]. Importantly, riociguat is approved to treat pulmonary arterial and chronic thromboembolic pulmonary hypertension [43]. sGC stimulators continue to undergo intensive preclinical and clinical testing for a variety of indications, including heart failure [44]. Recently, vericiguat successfully completed a phase 3 pivotal study in patients with chronic heart failure with reduced ejection fraction [45]. The purpose of the present study was to determine if the sGC stimulator BAY41-8543, the prototype of riociguat with the same mechanism of action, could mitigate skeletal muscle and cardio-respiratory dysfunction in the mdx4cv mouse model of DMD.

## Results

### 1. Increased sGC activity in mdx mice

To evaluate the potential of sGC as a novel drug target in DMD, we first investigated sGC activity, expression and localization in the skeletal muscles of the mdx4cv mouse model of DMD. Unexpectedly, relative to wild type controls, baseline sGC activity was 60 %, 141 % and 274 % higher in mdx gastrocnemius, tibialis anterior and diaphragm muscles (Figure 1a, d, e), respectively. Increased baseline sGC activity led to higher absolute sGC activities in mdx muscles in response to acute agonist treatment with the nitric oxide donor DETANO, sGC stimulator BAY41-2272 and sGC activator cinaciguat (Figure 1b, d, e). Note that BAY41-2272 was used in in vitro testing of sGC activity only and BAY41-8543 was used to treat mdx mice. The increase in sGC activity in response to agonists was similar between wild type and untreated and treated mdx mice (Figure 1c).

**Figure 1.**
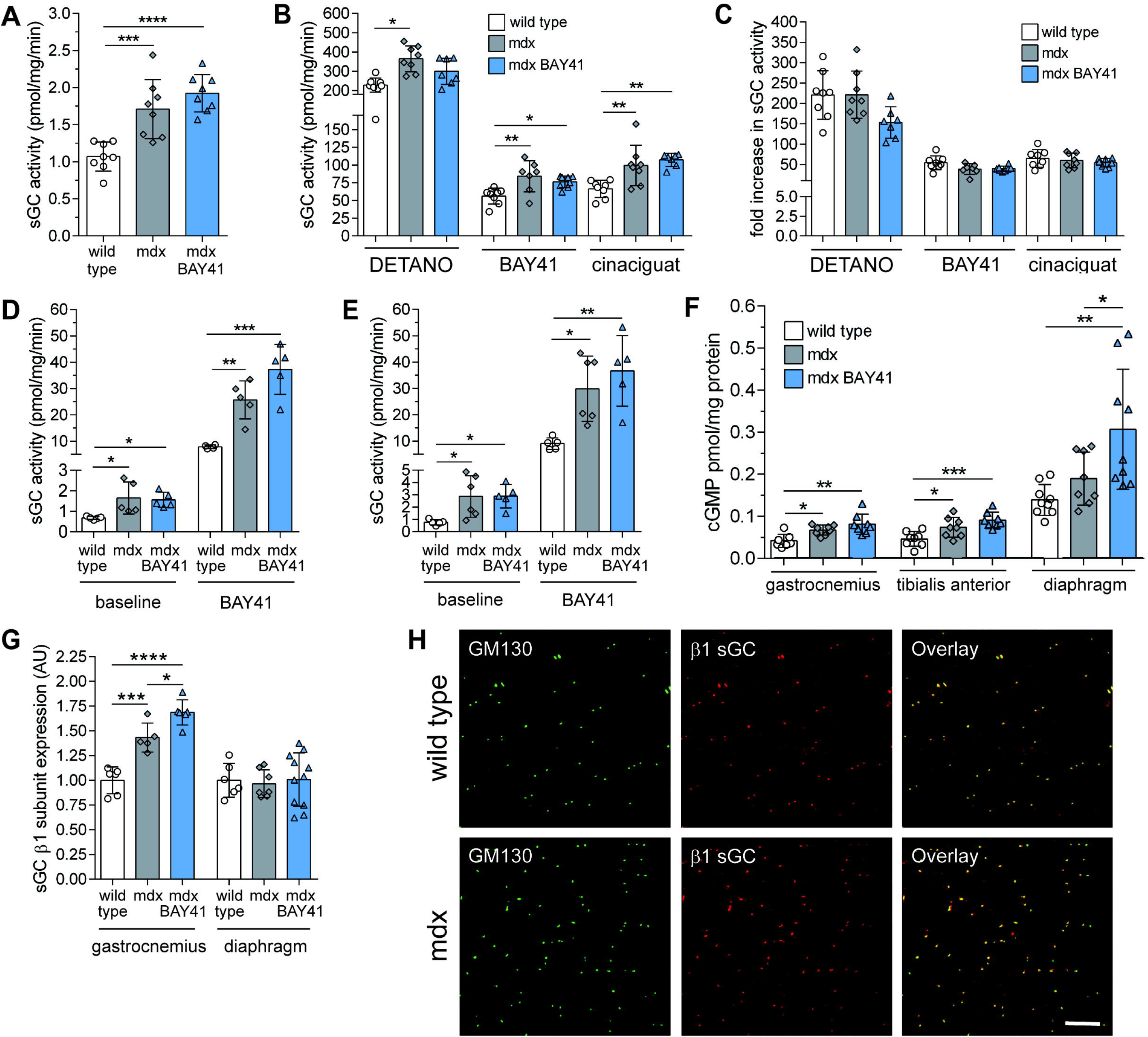
Nitric oxide-sensitive sGC activity is increased in mdx skeletal muscles. (A) Baseline sGC activity in resting gastrocnemius muscle from 15-week-old wild type, untreated and treated (3 months of BAY41-8543) mdx mice. (B) sGC activity in gastrocnemius muscles from wild type, untreated and treated mdx mice in response to acute exposure to functionally distinct agonist classes including nitric oxide donor DETANONOate (DETANO), heme-dependent sGC stimulator BAY41-2272 and heme independent sGC activator cinaciguat. (C) Fold increase in sGC activity over basal in response to different agonists in gastrocnemius muscles. (D) sGC activity at baseline and in response to acute BAY41-2272 exposure in tibialis anterior muscles. (E) sGC activity at baseline and in response to acute BAY41-2272 exposure in diaphragm muscles. (F) Steady state cGMP levels. (G) Expression of β1 sGC subunit, common to all sGC enzymes, determined by Western immunoblot. (H) Confocal micrographs of single myofibers from the gastrocnemius immuno-labeled with antibodies against cis-Golgi protein GM130 and β1 sGC. Scale bar 10 microns. * p < 0.05, ** p < 0.01, *** p < 0.001, **** p < 0.0001 by one factor ANOVA with Tukey’s multiple comparison test.

Accordingly, increased sGC activity was associated with a 60 % in the steady state cGMP pool in mdx gastrocnemius and tibialis anterior muscles and trended 35 % higher in the mdx diaphragm compared to controls (Figure 1f). In contrast to skeletal muscles, baseline cardiac sGC activity was similar between wild type and mdx mice (5.44 ± 0.87 and 5.18 ± 1.11 pmol/mg/min, respectively) and in response to DETANO or BAY41-2272 (not shown). These data indicate differences in NO-sGC-cGMP signaling between skeletal and cardiac muscle in mdx mice.

Increased sGC activity was accompanied by increased β1 sGC subunit expression in the gastrocnemius, but not diaphragm muscle (Figure 1g, Supplementary Figure 1a and c). α1 sGC subunit was unaffected thus increased α1β1 sGC may not be driving increased activity (Supplementary Figure 1b, d and e). Given the disruption of nNOS localization in mdx muscles, we next evaluated sGC localization. sGC localizes near the cis-Golgi complex in a “beads on a string” distribution in wild type myofibers (Figure 1h) [12]. Although the stereotypical localization of the cis-Golgi (marked by GM130) and sGC is disrupted in mdx myofibers, the close proximity of sGC to the Golgi complex was preserved (Figure 1h). Taken together, these findings indicate that in mdx skeletal muscles sGC activity is upregulated culminating in a larger steady state GMP pool. In some muscles increased sGC enzyme expression may be driving the increase in sGC activity.

### 2. BAY41-8543 treatment regimen

Given that upregulation of sGC activity could compensate for the 80 % reduction in nNOSμ activity in mdx skeletal muscles, we hypothesized that increasing sGC activity further would reduce dystrophic pathology. To test this hypothesis, we treated three weeks old mdx mice for three months with the sGC stimulator BAY41-8543 compounded into chow. This dosing regimen was efficacious in producing a ∼four-fold increase in lung cGMP (Supplementary Figure 2a). BAY41-8543 was well-tolerated with no effects on body weight despite an 18 % increase in food intake consistent with a thermogenic activity of cGMP in muscle and brown adipose tissue (Supplementary Figure 2b, c and d) [32, 46, 47]. This dose does not affect systemic blood pressure or heart rate in control C57BL6/J mice and had a minor blood pressure lowering effect in mdx mice during the active night cycle (Supplementary Figure 2e and f) [48].

**Figure 2.**
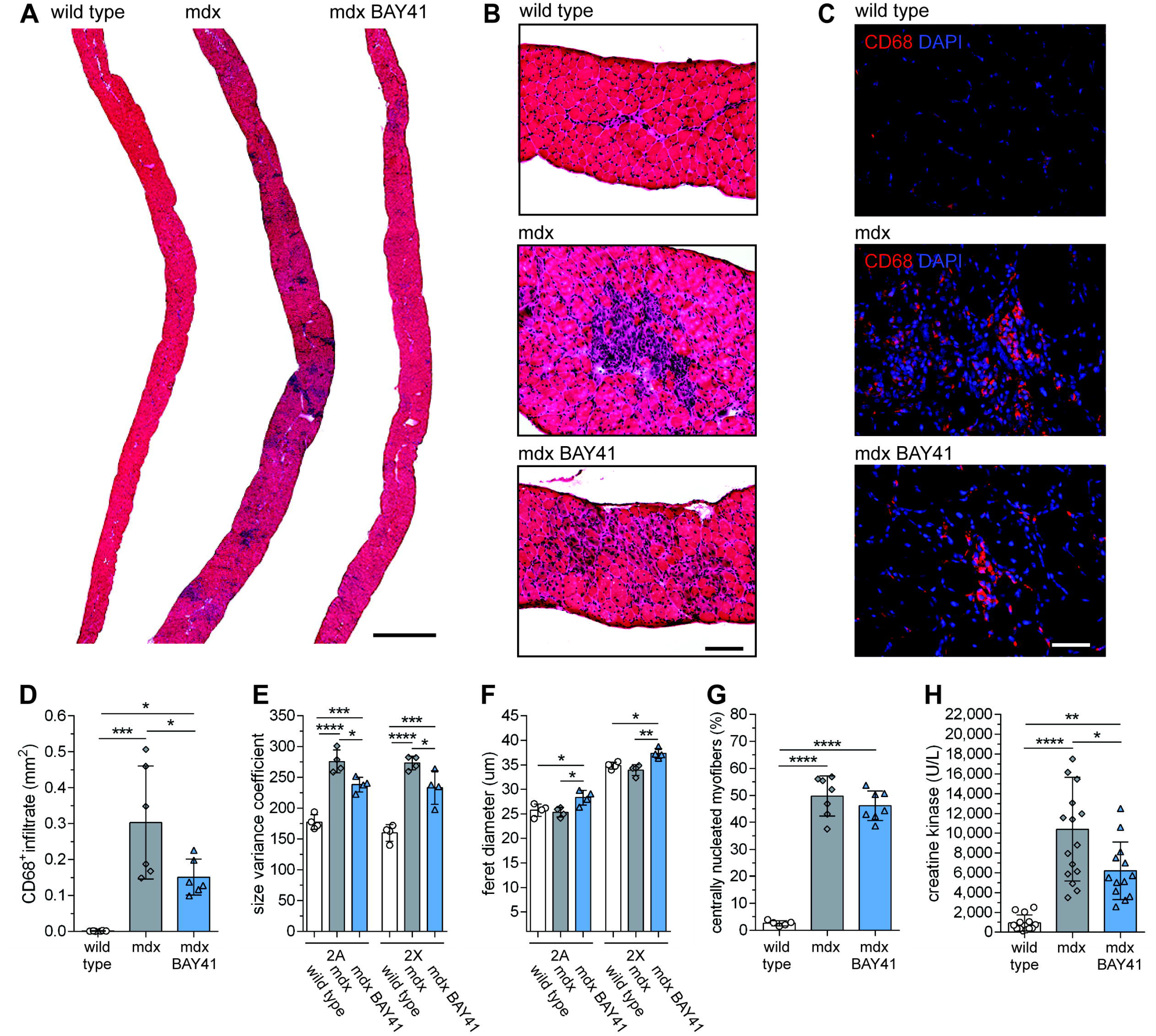
sGC stimulation reduces respiratory muscle inflammation and damage. (A) Diaphragm muscles from 15-week-old control, mdx and BAY41-8543 treated mice stained with hematoxylin and eosin to label nuclei and the cytoplasm, respectively. Scale bar 1 mm. (B) High magnification images in A highlighting inflamed area in mdx muscles marked by the blue nuclei of infiltrating immune cells. (C) Wide field immunofluorescence images of diaphragm muscles immunolabeled with anti-CD68 antibody (monocyte/macrophage marker-red) and counterstained with nuclear dye DAPI (blue). Scale bar 100 microns. (D) Quantitation of the area of CD68^+^ monocyte/macrophage infiltrate in diaphragm muscle sections. (E) Calculation of type 2A and 2X diaphragm myofiber size variability using the variance coefficient. (F) Determination of type 2A and 2X diaphragm myofiber size using minimum ferets’ diameter. (G) Percentage of centrally nucleated diaphragm myofibers. (H) Circulating serum creatine kinase activity. * p < 0.05, ** p < 0.01, *** p < 0.001, **** p < 0.0001 by one factor ANOVA with Tukey’s multiple comparison test.

### 3. sGC stimulation mitigates mdx skeletal muscle damage and inflammation

We evaluated the impact of BAY41-8543 treatment on diaphragm respiratory muscle damage (Figure 2). As expected, hematoxylin and eosin stained mdx diaphragm muscles showed substantial immune cell infiltration characterized by large accumulations of CD68 positive monocytes and macrophages (Figure 2a-d). BAY41-8543 treatment dramatically reduced CD68 positive immune cell infiltration (Figure 2a-d). Experimentalists blinded to treatment correctly identified treated hematoxylin and eosin stained mdx diaphragms 94 % of the time (n=8 per group) indicating that the anti-inflammatory effect of BAY41-8543 was clearly visually discernable.

Relative to wild type controls, mdx diaphragms also showed stereotypical increases in type 2 myofiber size variability and centrally nucleated myofibers (Figure 2e and g, respectively). BAY41-8543 significantly reduced type 2 myofiber size variability in mdx mice (Figure 2e), but did not affect central nucleation (Figure 2g). In addition, decreased inflammation in treated mdx mice was accompanied by myofiber growth indicated by increased feret diameter (Figure 2f). Importantly, serum creatine kinase activity in BAY41-8543 treated mdx mice was ∼40 % lower than untreated mdx mice consistent with treatment reducing global muscle damage (Figure 2h).

BAY41-8543 treatment had similar effects in mdx soleus muscles with reductions in immune cell infiltration clearly evident in treated mdx soleus muscles compared with untreated mdx controls (Figure 3a and b). BAY41-8543 had no impact on normalized mdx soleus muscle mass (Figure 3c), increased mdx myofiber size variability (Figure 3d) or the number of central nucleated myofibers (Figure 3f). However, as seen in the mdx diaphragm, BAY41-8543 also increased mdx type 2 myofiber feret diameter (compare Figure 3e to Figure 2f). Thus, the therapeutic benefits of BAY41-8543 were greater in the more affected mdx diaphragm. Collectively, these data indicate that BAY41-8543 treatment significantly reduced skeletal muscle inflammation and damage while promoting myofiber growth.

**Figure 3.**
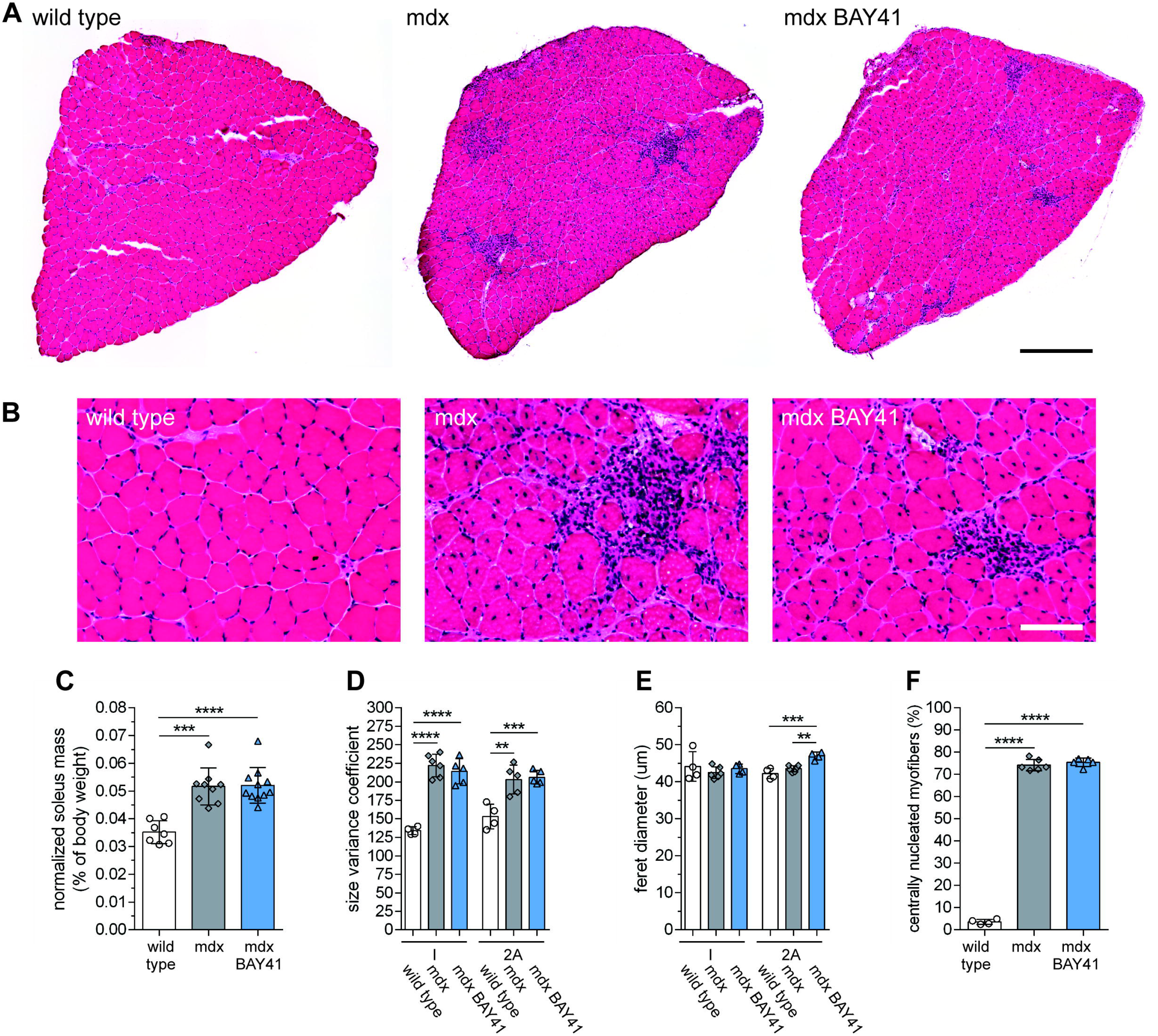
sGC stimulation reduces hind limb muscle inflammation and damage. (A) Soleus hind limb muscles from 15-week-old control, mdx and BAY41-8543 treated mice stained with hematoxylin and eosin to label nuclei and the cytoplasm, respectively. Scale bar 400 microns. (B) High magnification images of solei in A highlighting inflamed areas in mdx muscles marked by the blue nuclei of infiltrating immune cells. Scale bar 100 microns. (C) Impact of BAY41-8543 on soleus muscle mass as a percentage of body weight. (D) Type 2A and 2X soleus myofiber size variability using the size variance coefficient. (E) Minimum ferets’ diameter of type 2A and 2X soleus myofibers. (F) Percentage of centrally nucleated soleus myofibers. * p < 0.05, ** p < 0.01, *** p < 0.001, **** p < 0.0001 by one factor ANOVA with Tukey’s multiple comparison test.

To determine the strength of the anti-inflammatory effect of sGC stimulation, we tested if sGC stimulation could decrease peak macrophage inflammation in the muscles of 4-6 week old mdx mice. We quantitated muscle macrophage densities by flow cytometry and circulating cytokines and chemokines by Bio-Plex immunoassay in five weeks old mdx mice that received BAY41-8543 in their diet for only two weeks (Figure 4). As expected, 5 weeks old mdx muscles exhibited substantial monocyte and macrophage infiltration relative to wild type controls (Figure 4a-c) associated with elevated serum activity of the muscle damage marker creatine kinase (Figure 4d). In agreement with immunohistochemical analysis (Figure 2c), sGC stimulation significantly reduced the number of skeletal muscle CD45^+^/ CD11b^+^/ CD68^+^ monocytes and CD45^+^/ CD11b^+^/ CD68^+^/ F4-80^+^ macrophages by ∼62 % (Figure 4a and b). Treatment significantly lowered the number of macrophages per mg of mdx muscle by 61 % (Figure 4c). Reduced macrophage densities in treated mdx mice were accompanied by a 54 % reduction of creatine kinase activity indicating reduced global muscle damage (Figure 4d).

**Figure 4.**
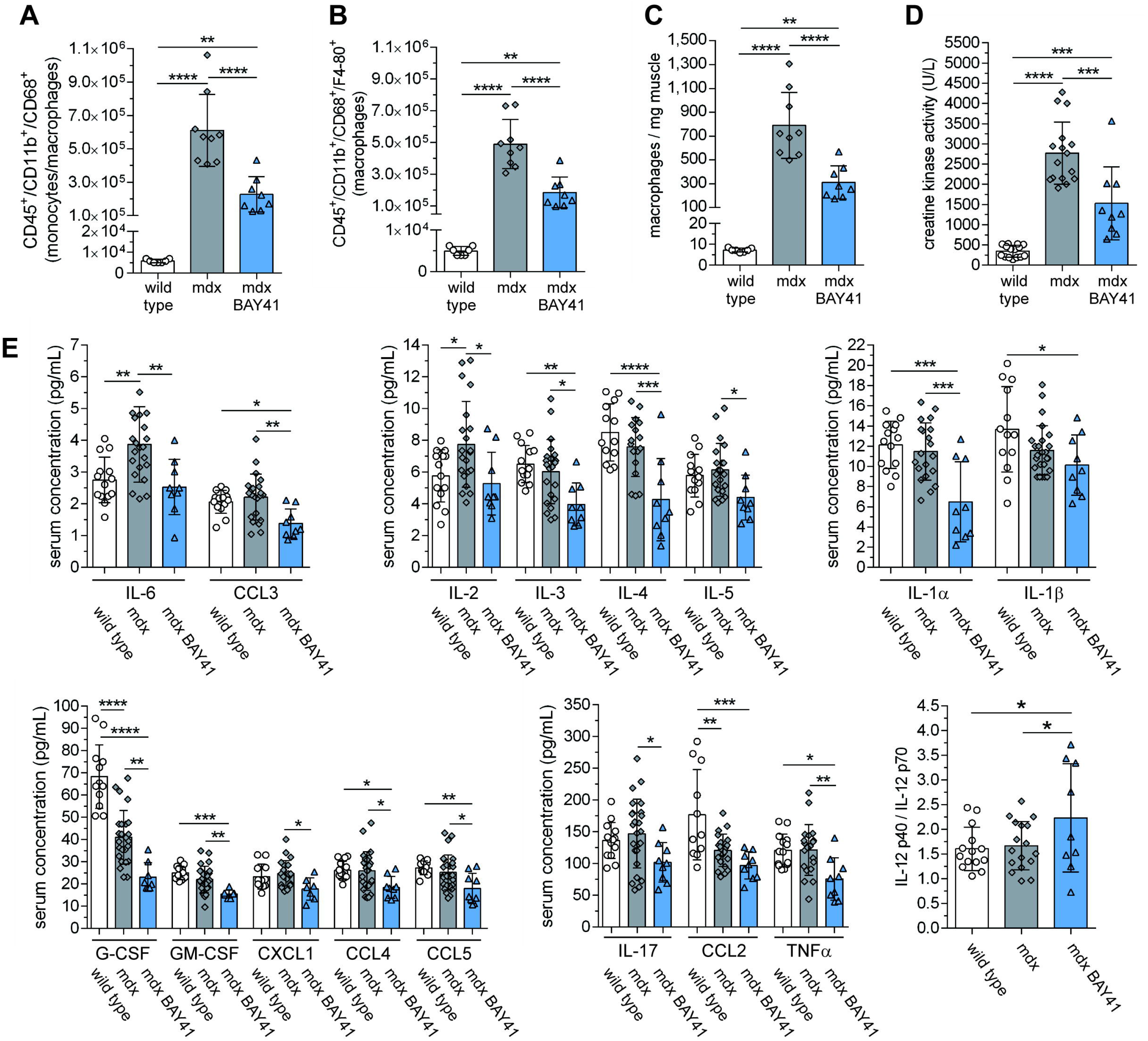
sGC stimulation reduces skeletal muscle macrophage densities and circulating pro-inflammatory cytokines and chemokines in mdx mice. (A) Flow cytometry quantitation of CD45^+^/CD11b^+^/ CD68^+^ monocytes/macrophages in the skeletal muscles of 5-week-old wild type, untreated mdx mice or mdx mice treated for 2 weeks with BAY41-8543. (B) Flow cytometry quantitation of CD45^+^/CD11b^+^/ CD68^+^/F4-80^+^ macrophages. (C) The number of macrophages normalized to skeletal muscle mass. (D) Serum creatine kinase activity in 5-week-old wild type, untreated mdx mice or mdx mice treated for 2 weeks with BAY41-8543. (E) Circulating concentrations of pro-inflammatory cytokines (IL-1 to 6, 17, G-CSF, GM-CSF and TNFα) and chemokines (CCL2-5 and CXCL1). 1-way ANOVA with either Holm-Sidak’s (A-D) or Tukey’s (E) multiple comparison tests. * p < 0.05, ** p < 0.01, *** p < 0.001, **** p < 0.0001.

We also assessed the impact of sGC stimulation on a battery of circulating pro-inflammatory mediators including IL-6 which can exacerbate dystrophic pathogenesis (Figure 4e) [49]. Interleukin 6 (IL-6) and 2 (IL-2) were significantly higher in mdx mice than in wild type controls. sGC stimulation of mdx mice reduced both IL-6 and IL-2 levels to wild type levels (Figure 4e). The circulating levels of most pro-inflammatory mediators in treated mdx mice were lower than untreated mdx or wild type controls with the exception of IL-1β and CCL2 (Figure 4e). Finally, sGC stimulator treatment with BAY41-8543 improved the ratio of IL-12 p40/p70 subunits (Figure 4e). Taken together, these data indicate that sGC stimulation can attenuate the immune response both locally in skeletal muscle by reducing muscle macrophage concentrations and globally by normalizing circulating serum IL-6 and IL-2 concentrations in mdx mice.

Next, we investigated if improvements in dystrophic muscle inflammation, damage and growth in sGC stimulator treated mdx mice were associated with reduced fibrosis (Figure 5, Supplementary Figure 3). Myofibers lacking dystrophin displayed excessive stereotypic endomysial fibronectin deposition compared with wild type controls (Figure 5a). Fibronectin also strongly marked fibrotic inflammatory lesions (Figure 5a. third panel-yellow box, Supplementary Figure 3a). sGC stimulation qualitatively reduced fibronectin deposition around mdx myofibers to a thickness comparable to wild type and reduced the size of fibronectin-positive fibrotic lesions (Figure 5a. Supplementary Figure 3a). Despite fibronectin remodeling to a distribution comparable to wild type, we observed no change in fibronectin protein expression (Figure 5c).

**Figure 5.**
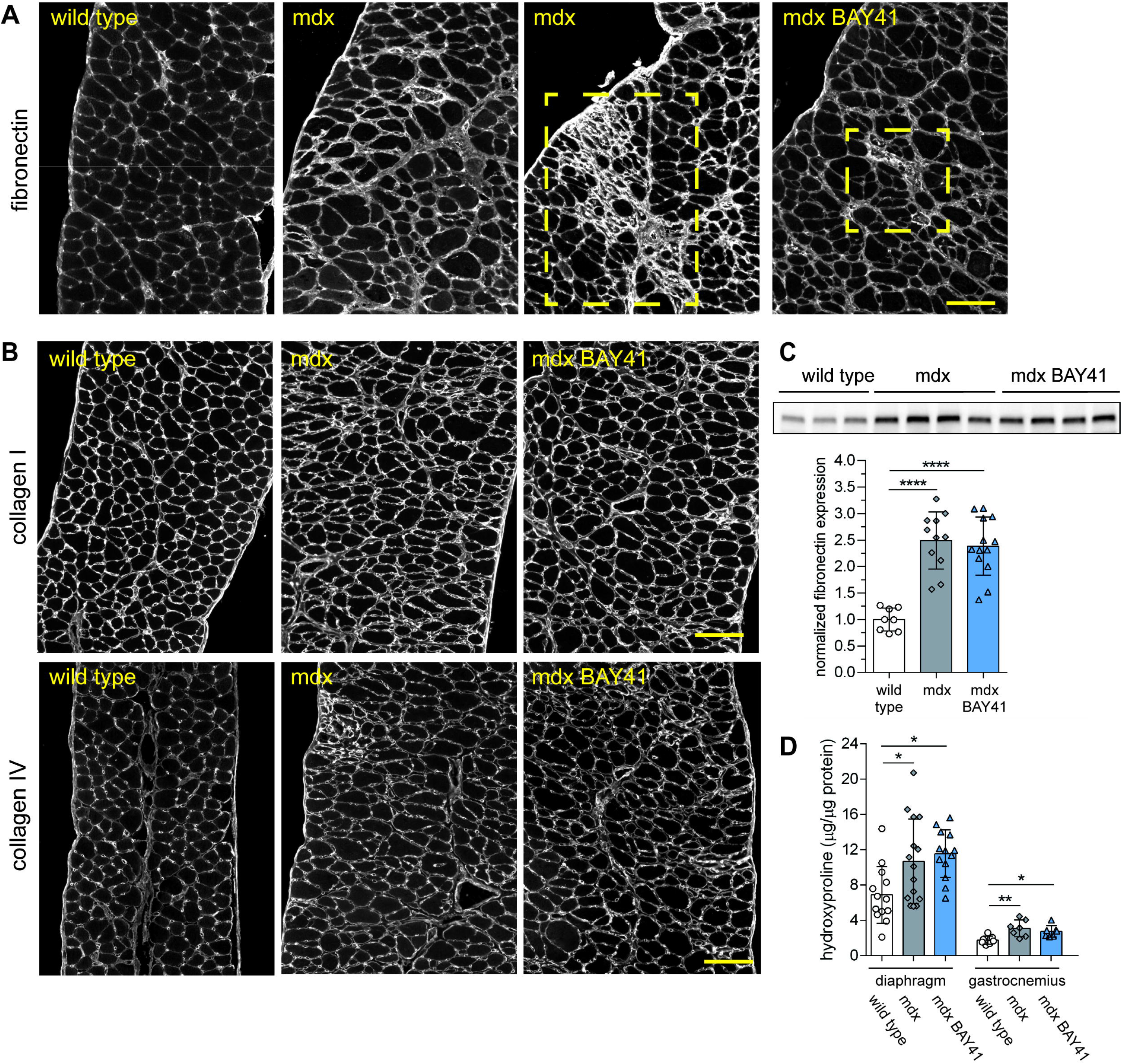
sGC stimulation reduces fibronectin positive lesions and promotes remodeling of fibronectin to a wild type distribution in the mdx diaphragm. (A) Representative wide field immunofluorescence images of diaphragm muscle cryosections from wild type, untreated and BAY41-8543-treated mdx mice immunolabeled with anti-fibronectin antibodies. (B) Representative wide field immunofluorescence images of diaphragm sections immunolabeled with anti-collagen I or anti-collagen IV antibodies (C) Representative Western immunoblot and densitometric quantitation of fibronectin protein expression. (D) Quantitation of hydroxyproline content as a measure of total collagen content. * p < 0.05, ** p < 0.01, *** p < 0.001, **** p < 0.0001 by one-factor ANOVA with Tukey’s multiple comparison test. Scale bar 100 microns.

sGC stimulation had no impact on excessive collagen I, III and IV deposition or lesions in mdx diaphragm muscle (Figure 5b and Supplementary Figure 3b). While hydroxyproline levels (a marker of bulk collagen) were elevated in both mdx diaphragm and gastrocnemius muscles compared with controls (Figure 5d), hydroxyproline levels were unaffected by BAY41-8543 treatment (Figure 5d). In agreement, elevated mdx muscle α-smooth actin protein expression, a marker of pro-fibrotic collagen-secreting myofibroblasts, was unaffected by sGC stimulation (Supplementary Figure 3c). Collectively, these data suggest that sGC stimulation had little effect on aberrant collagen deposition, but may remodel fibronectin-based fibrosis in mdx mice.

Then, we explored two possible mechanisms to further explain the myoprotective action of sGC stimulation. Increases in skeletal muscle NO-cGMP signaling can drive mitochondrial biogenesis [50]. Therefore, we first investigated if sGC stimulator mediated increases in cGMP could increase mitochondrial content in mdx muscle. sGC stimulation did not affect the expression of subunits of complex I, II, III, IV or V in mdx muscle (Supplementary Figure 4a) or the expression of mitochondrial voltage-dependent anion channel 1 (VDAC1) in mdx muscles (Supplementary Figure 4b). These findings argue that the myoprotective effects of sGC stimulation are not due to an increase in mitochondrial content.

Second, it is well established that increases in expression of the dystrophin homologue utrophin mitigate dystrophic pathology in mdx mice [51, 52]. Therefore, we investigated if sGC stimulation could increase utrophin protein expression. As expected, mdx diaphragm muscles expressed more utrophin than wild type controls (Supplementary Figure 4c). However, sGC stimulation had no effect on elevated utrophin expression in mdx mice arguing that the myoprotective effects of sGC stimulation are not due to a further increase in utrophin.

### 4. Long term sGC stimulation improves mdx skeletal muscle contractile performance

To determine if improved dystrophic muscle pathology translated to improved muscle function, we investigated if sGC stimulation could improve the contractile performance of mdx tibialis anterior muscles in situ (Figure 6). sGC stimulation had no impact on maximum tetanic force (Figure 6a) and did not affect specific force deficits at stimulation frequencies between 10 and 200 Hz (Figure 6b). Similarly, normalized force output-frequency curves were similar between untreated and treated mdx muscle (Figure 6c). In addition, sGC stimulation had no impact on the isometric twitch properties of mdx muscles (Supplementary Figure 5). Lastly, mdx tibialis muscles showed significant stereotypic deficits in force output during a series of repeated lengthening contractions compared to wild type (Figure 6d). However, BAY41-8543 treated mice showed moderately higher force output relative to untreated mdx controls (Figure 6d). This finding suggesting treatment improved resistance to lengthening contraction induced injury.

**Figure 6.**
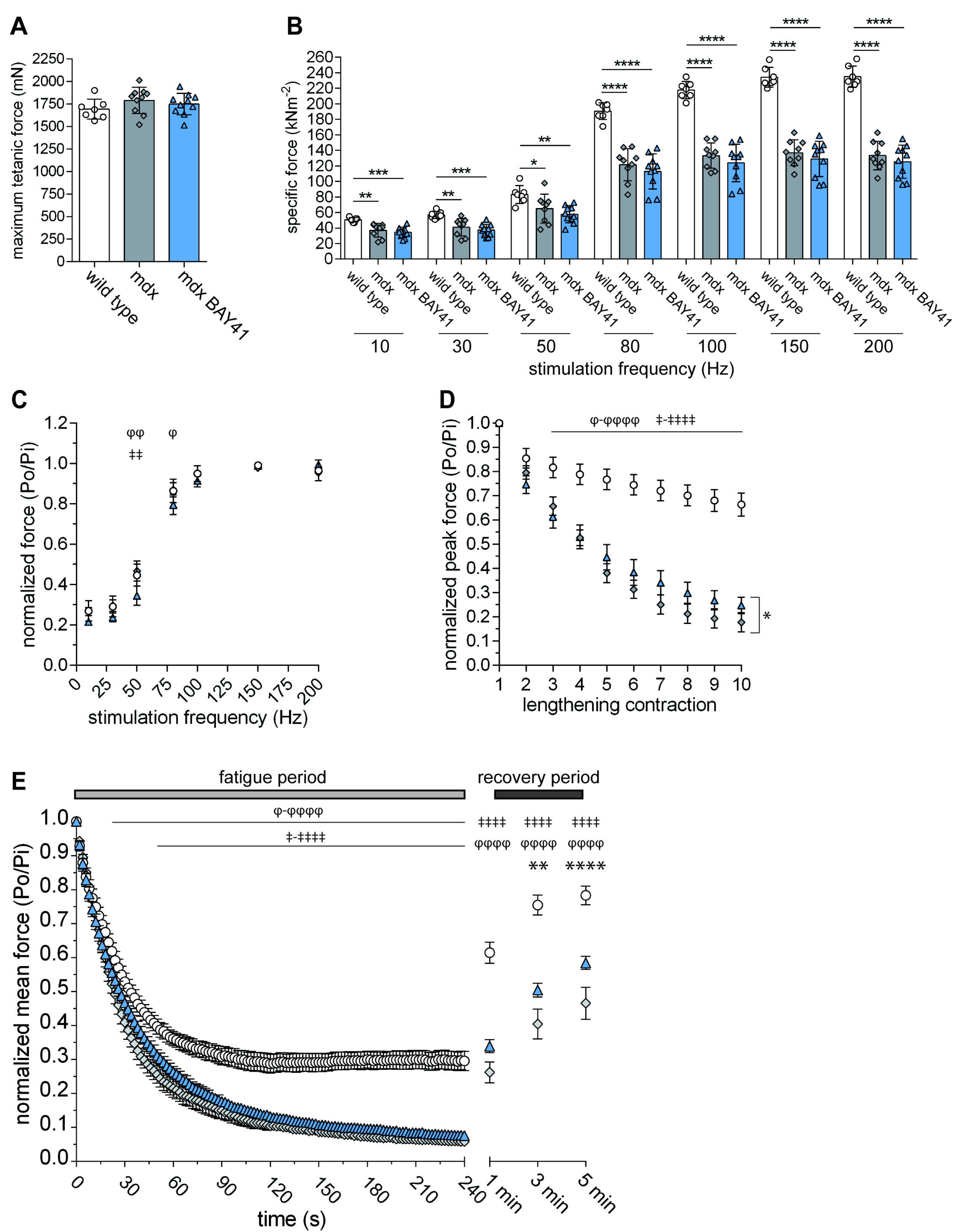
sGC stimulation improves mdx muscle resistance to contraction-induced fatigue. In situ analysis of isometric and eccentric (lengthening) tetanic force output of tibialis anterior muscles from wild type and untreated or 12-week BAY41-8543 treated mdx mice. (A) Maximum tetanic isometric force in tibialis anterior muscles from wild type and untreated or BAY41-8543 treated mdx mice. (B) Specific force output at submaximal and maximal peroneal nerve stimulation frequencies. * p < 0.05, ** p < 0.01, *** p < 0.001, **** p < 0.0001 by one factor ANOVA with Tukey’s multiple comparison correction. (C) Normalized force-frequency curves. (D) Normalized peak force output during 10 sequential lengthening (20 % stretch beyond optimum muscle length) contractions. (E) In situ analysis of the resistance of the tibialis anterior muscle to contraction-induced fatigue. Mean ± SEM force output (Po) normalized to initial mean force (Pi) during a four minute contraction-induced fatigue period and 5 minute recovery period. For C-E, wild type, mdx and mdx BAY41-8543 group sizes are 7, 9 and 10, respectively. Data presented as mean ± standard error. φ represents wild type versus mdx, ‡ represents wild type versus mdx BAY41-8543 and * represents mdx versus mdx BAY41-8543. Data analyzed by two factor ANOVA with Holm-Sidak correction for multiple comparisons where φ-φ φ φ φ represents p < 0.05-0.0001. ‡-‡‡‡‡ represents p < 0.05-0.0001. * represents P < 0.05, ** represents p < 0.01 and **** p < 0.0001.

The benefits of sGC stimulation were not restricted to improved eccentric contraction performance because post exercise contraction-induced fatigue was also improved by treatment (Figure 6e). During a four-minute fatigue protocol, mdx muscles showed significantly greater force deficits relative to wild type controls. During this period, stimulator treatment did not significantly impact contraction-induced fatigue resistance in mdx muscle (Figure 6e). However, treated muscles showed increased forced output during the recovery period indicating improved post exercise force recovery. Taken together, these findings suggest that sGC stimulation improves both the resistance of mdx muscle to eccentric contraction induced injury and recovery from contraction-induced fatigue.

### 5. Long term sGC stimulation improves respiration in mdx mice

To determine if sGC stimulation could benefit breathing capacity in dystrophic mice, we used non-invasive ultrasonography and whole-body plethysmography to determine the performance of the diaphragm respiratory muscle and respiratory capacity, respectively (Figure 7). As expected, the thoracic displacement of the mdx diaphragm was significantly reduced by 36 % in mdx mice compared with wild type controls (Figure 7a). However, sGC stimulation significantly improved diaphragm displacement by almost 30 % in mdx mice relative to untreated mdx controls (Figure 7a). However, sGC stimulation had no significant impact on deficits in mdx respiration frequency (Figure 7b), tidal volume (Figure 7c) or minute volume (Figure 7d). However, compared to both untreated mdx and wild type controls, treated mdx mice showed a significantly lower rejected breath index indicating reduced breathing irregularities (Figure 7e).

**Figure 7.**
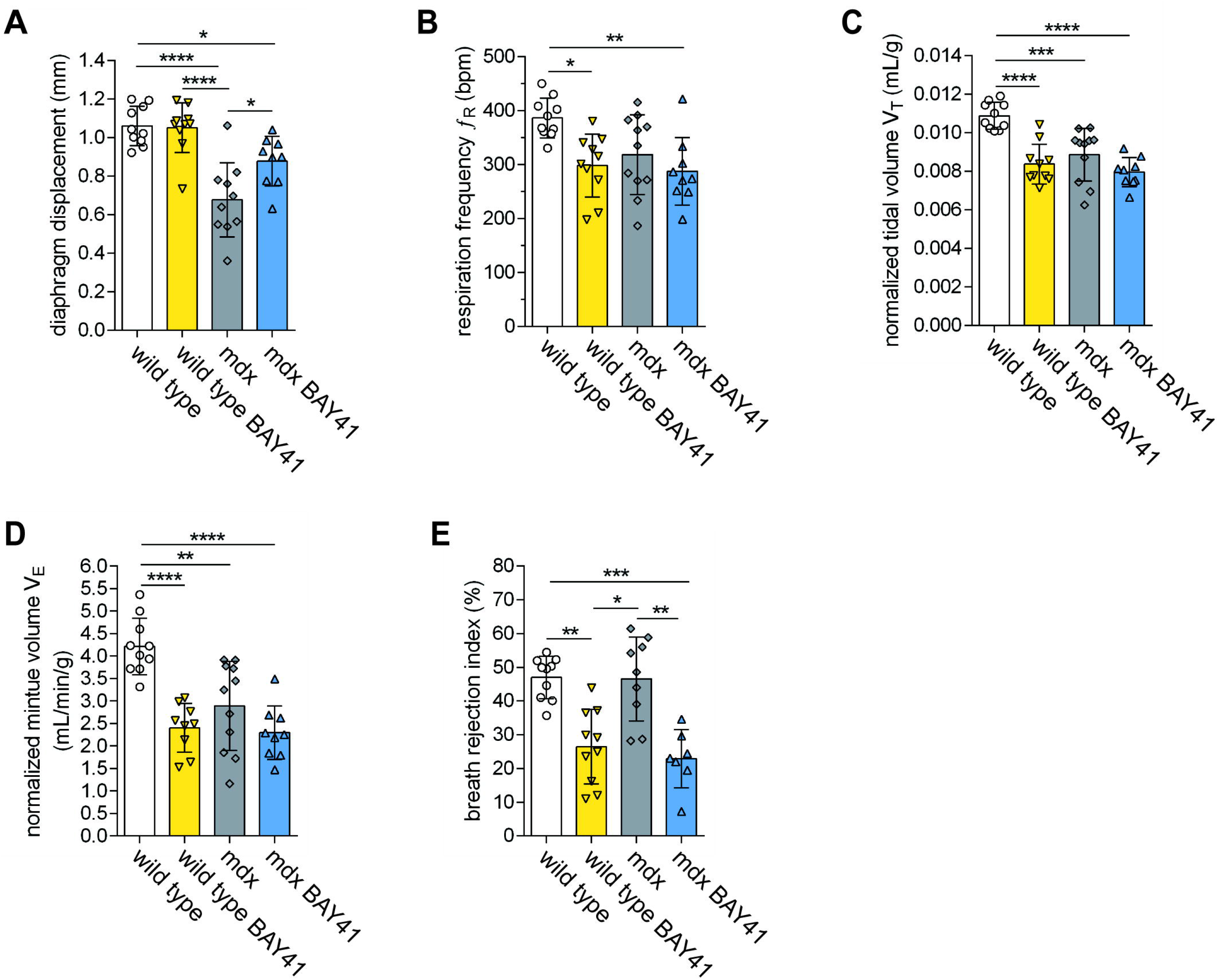
sGC stimulation reduces *in vivo* diaphragm dysfunction and breathing irregularities. (A) *In vivo* high-resolution ultrasonography analysis of the impact of 16-17 weeks of BAY41-8543 on diaphragm respiratory muscle displacement within the thoracic cavity of 19-20 week old untreated and BAY41-8543-treated wild type and mdx mice. (B-E) Whole body plethysmography analysis of the impact of BAY41-8543 on pulmonary function in 19-20 week old conscious and unrestrained untreated and BAY41-8543-treated wild type and mdx mice. (B) Respiratory rates. (C) Body mass-normalized tidal volumes. (D) Body mass-normalized minute volumes. (E) Impact of BAY41-8543 on the fraction of breaths rejected (breath rejection index or Rpef) for analysis. * p < 0.05, ** p < 0.01, *** p < 0.001, **** p < 0.0001 by one-factor ANOVA with Tukey’s post-hoc correction for multiple comparisons.

We also determined the impact of BAY41-8543 on respiratory muscle and breathing function in wild type mice. sGC stimulation had no impact on wild type diaphragm displacement (Figure 7a). However, compared with untreated wild type controls, sGC stimulated wild type mice exhibited quieter and more regular breathing with significant decreases in respiration frequency (Figure 7b), tidal volume (Figure 7c), minute volume (Figure 7d) and breath rejection index (Figure 7e). The breathing patterns of treated wild type mice resembled those of untreated mdx mice.

Compared with untreated wild type mice, sGC stimulated wild type mice also exhibited significant increases in inspiratory and expiratory times and total breathing cycle time (Supplementary Figure 6a-c). Importantly, altered respiration in treated wild type was similar to untreated and treated mdx mice in most parameters tested except breath rejection index which was improved by sGC treatment in both wild type and mdx mice (Figure 7e). In addition, BAY41-8543 treated wild type mice exhibited significant decreases in peak inspiratory flow, peak expiratory flow and expiratory flow at 50 % expired volume compared with untreated wild type controls (Supplementary Figure 6d-f). Taken together these data suggest that increasing sGC activity in wild type mice is sufficient to decrease or quieten breathing comparable to that in mdx mice without impacting diaphragm respiratory muscle displacement. It is important to note that cardiac function in these mice was comparable to wild type at this age (not shown). These findings suggest that sGC stimulator treatment improves both respiratory muscle displacement and breathing quality in mdx mice.

### 6. sGC stimulation mitigates diastolic dysfunction and fibrosis in aged mdx hearts

We then determined the effect of two months of sGC stimulation on cardiomyopathy in 15-month-old mdx mice (Figure 8). Steady state cGMP levels were similar between wild type and mdx hearts (Figure 8a). However, sGC stimulator treatment significantly increased cGMP levels compared to untreated groups (Figure 8a). Heart rate was similar between all groups (Figure 8b). Stroke volume and cardiac output were lower in mdx mice than in wild type controls, but unaffected by treatment (Figure 8c and 8d). The worsening of left ventricle end systolic volume (LVESV) and diastolic volume (LVEDV) in mdx mice relative to wild type controls was also unaffected by treatment (Figure 8e-f). sGC stimulator treatment also did not impact other indices of systolic dysfunction in mdx mice including ejection fraction % and fractional shortening % (Figure 8g, Supplementary Figure 7a). Taken together, these data suggest negligible impact of sGC stimulation on systolic dysfunction in mdx mice.

**Figure 8.**
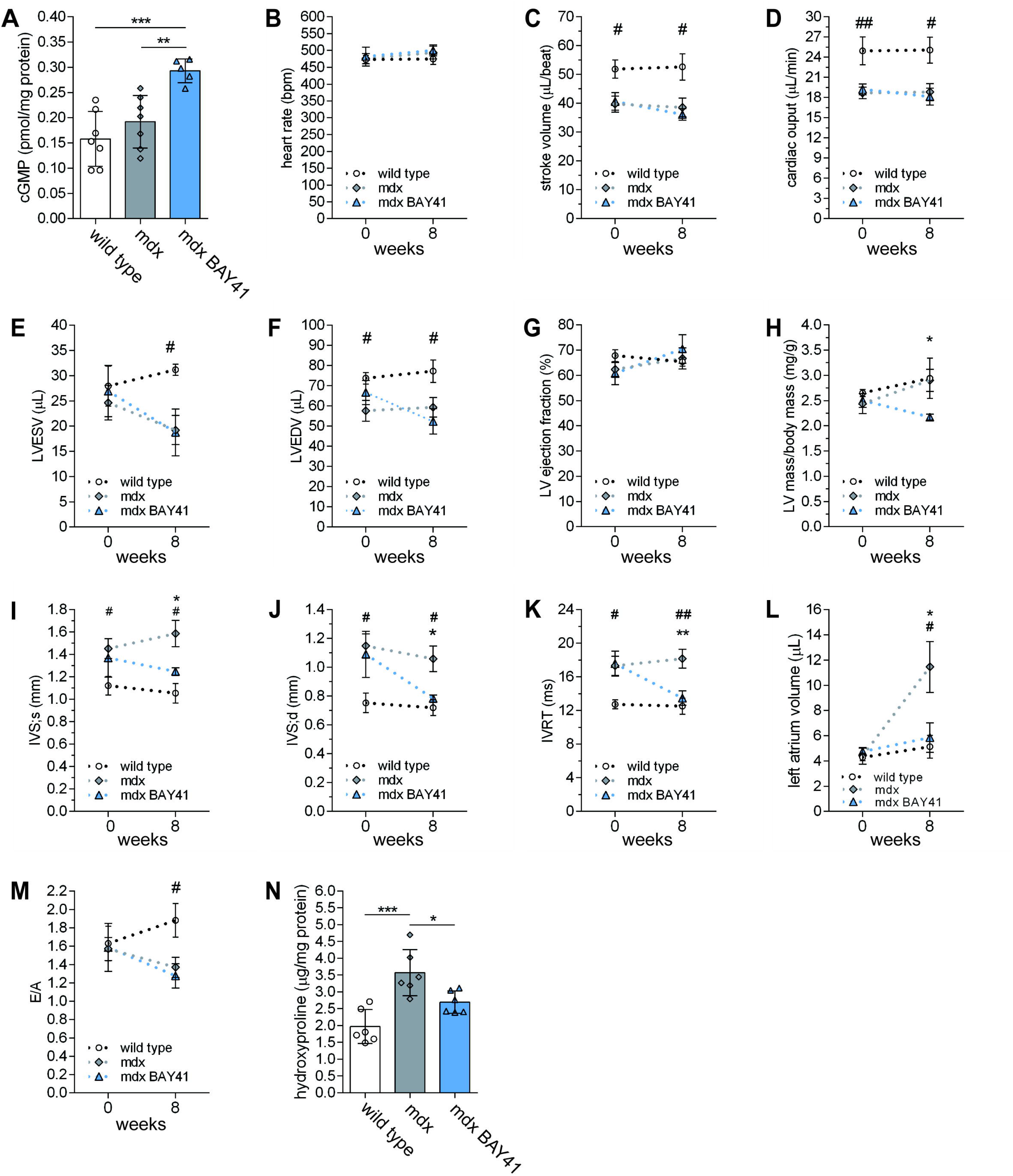
sGC stimulation ameliorates diastolic dysfunction in aged mdx hearts. Impact of 8 weeks of BAY41-8543 on cardiomyopathy in mdx mice starting at 15 months of age. (A) Steady state cardiac cGMP concentrations in wild type and untreated and BAY41-8543 treated mice. (B-D) Impact of BAY41-8543 on left ventricle (LV) systolic function and pathogenic chamber remodeling in mdx mice determined by prospective short axis M mode trans-thoracic echocardiography at baseline and 8 weeks after treatment. (B) Heart rate. (C) Stroke volume. (D) Cardiac output. (E) LV end systolic volume (LVESV). (F) LV end systolic volume (LVEDV). (G) LV ejection fraction. (H) Corrected LV mass normalized to body weight. (I) Interventricular septum thickness at systole. (J) Interventricular septum thickness at diastole. (K-M) Impact of BAY41-8543 on diastolic function determined by pulsed wave Doppler mode and by parasternal long axis B-mode (four chamber) analyses. (K) Isovolumic relaxation time. (L) Calculated left atrium volume. (M) Early (E) to late (A) transmitral blood flows. (N) Cardiac hydroxyproline content as a measure of fibrotic collagen deposition. Group sizes, 5-7, 8 and 6 for wild type, mdx and BAY41-8543-treated mdx, respectively. B-M, n=5-8, mean ± SEM, unpaired t-test where # and ## represents p < 0.05 and p < 0.01 for wild type versus mdx, respectively. * represents p < 0.05 for mdx versus mdx BAY41-8543. A and N, * p < 0.05, ** p < 0.01, *** p < 0.001, **** p < 0.0001 by one-factor ANOVA with Tukey’s post-hoc correction for multiple comparisons. N-One way ANOVA with Tukey’s correction for multiple comparisons.

In contrast, sGC stimulator treatment reduced hypertrophy of the interventricular septum in systole (IVS;s) in mdx mice and normalized IVS wall thicknesses in diastole (IVS;d) to wild type (Figure 8i and j). BAY41-8543 treatment prevented increases in body weight normalized left ventricle mass observed in both wild type and mdx mice and in anterior wall thickness at systole (LVAW;s) (Figure 8h, Supplementary Figure 8b). However, other measures of cardiac wall thickness were similar between wild type and mdx mice and unaffected by treatment (Supplementary Figure 8c-g). Taken together these data suggest that sGC stimulation opposed deleterious hypertrophic cardiac muscle remodeling in mdx mice.

Mdx mice exhibit a prolonged isovolumic relaxation time (IVRT) compared with wild type controls, indicative of impaired myocardial relaxation (Figure 8k) [53]. Therefore, we investigated the impact of sGC stimulation on diastolic dysfunction in aged mdx mice. sGC stimulator treatment normalized IVRT in mdx mice. In addition, relative to wild type controls, mdx mice developed substantial enlargement of the left atrium, which was prevented by sGC stimulator treatment (Figure 8l). As expected, mdx mice also showed a decrease in E/A ratio relative to wild mice indicating diastolic dysfunction (Figure 8m). However, BAY41-8543 treatment had no impact on E/A ratio.

Finally, we investigated whether sGC stimulator treatment could reduce fibrosis in aged mdx heart as a mechanism contributing to improved myocardial relaxation. As expected, hydroxyproline, a marker of collagen, was increased 81 % in mdx hearts relative to wild type controls (Figure 8n). sGC stimulation reduced cardiac collagen content in mdx hearts by 25 % (Figure 8n). These findings suggest that sGC stimulator treatment reduces diastolic dysfunction in dystrophic hearts by opposing deleterious left atrium enlargement and fibrosis.

## Discussion

The major finding of this study is that oral administration of sGC stimulator BAY431-8543 substantially reduced skeletal and cardiac muscle dysfunction at critical stages of disease progression in mdx mice. Importantly, the benefits of sGC stimulation in mdx mice occurred against the background of increased NO sensitive sGC activity and without a major effect on blood pressure. The broad range of therapeutic benefits provided by sGC stimulation are consistent with a key role for impaired NO-cGMP signaling in exacerbating dystrophic pathogenesis and reinforce the attractiveness of NO-cGMP pathways as a clinical target in DMD.

A key therapeutic benefit of sGC stimulation was reduced inflammation. Consistent with reported anti-inflammatory and macrophage suppressing functions of nNOS in mdx muscles, sGC stimulation significantly reduced muscle monocyte and macrophage concentrations [21, 26]. Reductions in macrophages during peak inflammation were accompanied by normalized levels of cytokines, including IL-6, reported to exacerbate diaphragm weakness [49, 54, 55]. Accordingly, reductions in inflammation were associated with an improvement in *in vivo* diaphragm displacements and fewer breathing abnormalities. Collectively, these findings suggest that the absence of intact nNOS-sGC signaling axis exacerbates inflammation in dystrophin-deficient muscle.

The reductions in inflammation in BAY41-8543 treated mdx mice were associated with substantial reductions in skeletal muscle damage without a change in central nucleation, and with modest muscle growth and remodeling of fibronectin positive fibrotic lesions. These data suggest sGC stimulation promoted a beneficial rebalancing of the inflammatory response with muscle growth and repair. Improvements in hind limb skeletal muscle health were associated with decreased contraction-induced muscle fatigue and susceptibility to lengthening contraction induced muscle damage. Previous reports indicated a role for nNOS and sGC in post exercise force recovery; therefore, improved contraction-induced muscle fatigue in mdx mice may be specifically due to increased sGC activity alone [32].

It is worth noting that in our hands the susceptibility of mdx4cv mice to contraction induced fatigue resistance is substantially higher than that of the most studied C57BL/10ScSn-Dmd^mdx^/J model which we and others have shown exhibits similar fatigue resistance to wild type controls [56]. The difference in fatigue resistance between mdx4cv and C57BL/10ScSn-Dmd^mdx^/J models are unlikely to be technical since the same approach was used in both mdx models and may related to differences in mutation (exon 23 versus 53) or background strain. To the best of our knowledge, this is the first report of such a critical difference and provides a compelling argument that going forward the mdx4cv mice be used in studies of muscle fatigue in DMD.

BAY41-8543 treated mice not only showed improved hind limb muscle function but also exhibited improved diaphragm respiratory muscle health and function associated with increased breathing quality. Mechanistically, improved diaphragm *in vivo* function could be due to an early reduction in circulating IL-6 reported to promote diaphragm dysfunction in mdx mice [49, 54, 55]. Importantly, consistent with the importance of sGC in respiratory and pulmonary function, sGC stimulation in wild type control mice decreased respiratory frequency and decreased tidal and minute volumes to levels comparable to what was observed in mdx mice (Figure 7b-e). BAY41-8543 decreased breathing irregularities to the same extent in both wild type and mdx mice indicating that the change was not related to dystrophin deficiency. First, these intriguing findings from treated wild type controls reveal a novel role for sGC in modulating respiratory capacity and quality that warrants further investigation. Second, these findings suggest that increased sGC activity in mdx mice may promote diaphragm health and *in vivo* function while simultaneously reducing breathing irregularities providing a novel therapeutic avenue for alleviating respiratory dysfunction in DMD.

Many pre-clinical studies employing genetic and pharmacological approaches (e.g., phosphodiesterase 5 (PDE5) inhibitors to elevate NO-cGMP signaling have reported reduced cardiac dysfunction and fibrosis in dystrophin deficient animals [27, 53, 57, 58]. In agreement, we found that BAY41-8543 treatment reduced age-dependent diastolic dysfunction and cardiac fibrosis in aged mdx mice that manifest more severe cardiomyopathy. Two months of sGC stimulation attenuated IVS hypertrophy and prevented progressive age-dependent maladaptive enlargement of the left atrium (Figure 8). This finding is supported by a previous study showing that inhibiting cGMP breakdown with sildenafil decreases IVS hypertrophy in younger mdx mice [53]. Thus, current evidence suggests that raising cGMP, either by stimulating sGC or blocking PDE5 activity, can block maladaptive hypertrophy in mdx hearts.

The therapeutic effect of sGC stimulation on left atrial remodeling could be highly clinically relevant. Enlargement of the left atrium is commonly associated with increased incidence of atrial fibrillation, stroke risk and overall mortality after myocardial infarction [59]. Left atrial enlargement also increases the risk of death and hospitalization in individuals with dilated cardiomyopathy that develops in many DMD patients [60]. Thus sGC stimulation could be useful in preventing left atrial remodeling and its deleterious sequelae in DMD patients.

Improvements in diastolic function were associated with a marked reduction in cardiac collagen, suggesting a strong anti-fibrotic action of BAY41-8543 in the heart, which is supported by previous studies of sGC stimulator efficacy in heart failure models [61]. In contrast, we did not observe a reduction in mdx diaphragm fibrosis with treatment. These data suggest important differences in the role of NO-sGC signaling in the development of fibrosis between cardiac and diaphragm muscle. Nonetheless, taken together these findings support an important role for NO-sGC-cGMP signaling in opposing diastolic dysfunction and fibrosis in dystrophin-deficient hearts.

The significant increase in skeletal muscle sGC activity in mdx mice was an unexpected finding. Skeletal muscle sGC activity and agonist activation requires nNOS. Accordingly, sGC activity is substantially decreased in nNOS deficient mice and in DMD patients [32, 33]. Our data indicated that baseline sGC activity and steady state cGMP levels were substantially higher in skeletal muscle of mdx mice than in WT mice in part because of increased sGC enzyme expression (Figure 1). These findings suggest an unrecognized species-specific compensation for the loss of nNOS activity in dystrophin-deficient murine but not human skeletal muscles. Importantly, these data raise the exciting possibility that sGC stimulators could be even more effective or therapeutic in restoring NO-sGC-cGMP signaling in DMD patients than in mdx mice.

Previous attempts to increase NO-sGC-cGMP signaling in DMD primarily involved the use of phosphodiesterase 5 (PDE5) inhibitors to block cGMP breakdown [15]. Many studies showed that PDE5 inhibitors provided substantial therapeutic benefit in skeletal and cardiac muscles of animal models of DMD and alleviated functional ischemia in individuals with DMD or BMD [41, 53, 57, 62]. However, a lack of efficacy of PDE5 inhibitors in mitigating cardiomyopathy and the decline in ambulatory ability in individuals with DMD or BMD has dampened enthusiasm for their clinical use [40, 42, 63]. Reduced efficacy of PDE5 inhibitors may be due to several factors including reduced cGMP and decreased PDE5 target expression that both negate the effects of PDE5 inhibitors, muscle use patterns and also patient age and thus disease progression. In addition, sGC mediates most of the cGMP dependent functions of nNOS, while PDE5 mediates only a small subset of nNOS functions. Taken together, these studies suggest that PDE5 inhibition may not be an ideal strategy for restoring nNOS-cGMP signaling in individuals with DMD.

The present study provides context as to why several groups were able to demonstrate that PDE5 inhibitors effectively mitigated striated muscle dysfunction in mdx mouse models. Our data indicate that baseline sGC activity and steady state cGMP levels were elevated in mdx mice. In contrast, cGMP synthesis is decreased about 70% in DMD patients suggesting that there is low sGC activity and little cGMP to protect from PDE5 breakdown. In contrast, in mdx mice there is high sGC activity and a lot of cGMP to protect from PDE5 breakdown. Therefore, unlike sGC stimulators, PDE5 inhibitors would be expected to be more effective in dystrophin-deficient mice than in dystrophin-deficient humans.

sGC stimulators avoid many of the limitations of PDE5 inhibitors, they are expected to be therapeutically more effective than PDE5 inhibition [38]. Indeed, comparison of the findings from the present study with previous reports with PDE5 inhibitors suggest that sGC stimulation of cGMP synthesis results in a broader range of therapeutic benefits in dystrophic mice. For example, unlike PDE5 inhibitors, sGC stimulation significantly reduced serum creatine kinase activity and improved resistance to contraction-induced muscle fatigue and lengthening contraction-induced muscle damage [56]. In addition, sGC stimulation mitigated maladaptive atrial enlargement and cardiac fibrosis in aged mdx mice not previously observed with PDE5 inhibitors. Collectively, these findings suggest that sGC stimulation should be therapeutically more effective than PDE5 inhibition and intriguingly this may be more apparent in DMD patients than in mdx mice.

In summary, this study provides compelling preclinical evidence that sGC stimulation significantly mitigates clinically relevant skeletal and cardio-respiratory dysfunction in an mdx mouse model of DMD. In doing so, this study provides a new framework for understanding and addressing the defects in NO-cGMP signaling in DMD. This framework includes the key identification of increased sGC activity as a novel murine adaptation to the loss of nNOS, which will likely have significant implications for understanding and therapeutically addressing NO signaling deficits in animal models and individuals with DMD. For example, sGC stimulation could be used alone or in combination with several gene therapy approaches that do not address nNOS-cGMP dysfunction in skeletal or cardiac muscle.to improve their efficacy in DMD. Finally, this study provides the first evidence for sGC as a novel target in DMD and that repurposing of sGC stimulators may be an effective new therapeutic approach in DMD.

## Acknowledgements

This work was supported by Department of Defense award MD140021 to JMP.

## Methods

### Animals and Drug Dosing Regimen

#### Animals

All experimental procedures performed on mice were approved by the Institutional Animal Care and Use Committees of the University of Miami. Blood pressure experiments were performed at Bayer AG, according to the guidelines approved by the local animal welfare authorities for the German state of North-Rhine Westphalia (Landesamt für Natur, Umwelt und Verbraucherschutz Nordrhein-Westfalen). Male mdx^4cv^ mice (#002378, The Jackson Laboratory) that harbor a premature stop codon within exon 53 of the *DMD* gene that prevents dystrophin expression were used [64, 65]. Exon 53 is within a mutational hot spot region in DMD [66]. mdx^4cv^ mice have fewer dystrophin positive revertant myofibers and exhibit more progressive striated muscle pathology than mdx^ScSn^ mice. Strain (C57BL6/J), age and sex matched mice were used as controls.

#### Animal dosing

Litters of males, kept together to minimize social stress, were numbered and randomly assigned to control or soluble guanylate cyclase (sGC) stimulator BAY41-8543 chow diet starting at 3 weeks of age for 2 or 12 weeks, or starting at 15 months of age for 2 months for cardiac studies. BAY41-8543 (provided by Bayer AG) is a sGC stimulator tool compound for preclinical use which exhibits the same mechanism of action as other sGC stimulators like BAY 41-2272, riociguat, vericiguat, praliciguat and olinciguat with a similar pharmacological profile [36]. BAY41-8543 was premixed at 150 ppm into Teklad Global 18 % Protein Rodent diet (Envigo, Teklad Diets, WI). This dose approximates 10-15 mg/kg/day according to mouse body mass. Control chow was identical in every way except that it lacked BAY41-8543. Food eaten per day per mouse was determined by subtracting the food remaining in the cage feeder and on the cage floor from the total food provided at the beginning of the week then diving by 7 days and the number of mice in the cage.

### Biochemical assays

Skeletal muscle soluble guanylate cyclase (sGC) activity was measured in muscle homogenates exactly as described [32]. cGMP was measured by immunoassay (Cayman Chemical). sGC activity was expressed as picomoles of cGMP synthesized per milligram of protein per minute. Steady state cGMP concentrations were measured using a cGMP Direct ELISA Kit according to the manufacturer’s directions (Arbor Assays).

Serum creatine kinase activity was measured as a biomarker of muscle damage using the Stanbio™ CK Liqui-UV™ Test (Thermo Fisher Scientific) on a Biotek Synergy H1 microplate reader according to the manufacturer’s instructions. Total collagen concentrations in skeletal and heart muscles were measured with a hydroxyproline assay kit (Chondrex, #6017) on a Biotek Synergy H1 microplate reader according to the manufacturer’s directions. Circulating cytokines, chemokines and growth factors were quantitated by using a Bio-Plex Pro Mouse 23-plex Immunoassay and Bio-Plex Reader System (Bio-Rad).

### Western Immunoblotting

Western immunoblotting of skeletal muscles was performed as described [32]. Muscles were homogenized in buffer (2% sodium dodecyl sulfate, 50□mM Tris-HCl, pH 6.8) with protease and phosphatase inhibitors (Roche). Protein concentrations were determined by Bicinchoninic acid assay (Sigma-Aldrich) and 30 μg of protein per sample was electrophoresed by SDS PAGE on 4–20% Mini-PROTEAN TGX Stain-Free Precast Gels (Bio-Rad) and transferred to polyvinylidene fluoride membranes (Millipore). Equal lane loading was determined using the stain-free detection system and a Bio-Rad ChemiDoc MP imager. Protein expression was normalized to total transferred protein using Bio-Rad ImageLab 5.2.1 software. Membranes were blocked with 5% (w/v) skim milk in tris-buffered saline with Tween-20 for 1□hour at room temperature and incubated with primary antibodies overnight at 4°C.

Primary antibodies raised against sGCα1, (#G4280) and sGCβ1 (#G4405) were purchased from Sigma-Aldrich. Primary antibodies to fibronectin (#ab2413), α-smooth muscle actin (#ab5694), and VDAC1 (#15895) were purchased from Abcam. MitoProfile® total OXPHOS antibody cocktail (Abcam) was used to detect respiratory complex subunits NDUFB8 (Complex I or CI), SDHB (CII), UQCRC2 (CIII), MTCO1 (CIV), and ATP5A (CV). The anti-utrophin antibody (#MANCHO3 8A4) was purchased from Developmental Studies Hybridoma Bank, University of Iowa). Primary antibodies were detected with donkey horseradish peroxidase-conjugated secondary antibodies (Jackson ImmunoResearch Laboratories). Chemiluminescence was initiated using SuperSignal West Femto Maximum Sensitivity substrate (Thermo Fisher) and visualized on a Bio-Rad ChemiDoc MP imaging system.

### Immunohistochemistry and histological analyses

#### Single myofiber immunofluorescence labeling and confocal imaging

Gastrocnemius muscles were mechanically separated into small bundles or single myofibers with tweezers, then fixed in 2 % paraformaldehyde and immunolabeled with FITC-conjugated GM130 (#612008, BD Biosciences) and sGCβ1 (#G4405, Sigma-Aldrich) antibodies as described previously [67]. Confocal images were captured with a FluoView FV1000 confocal microscope (Olympus). Identical laser power, excitation, and emission capture settings were used to compare sGCβ1 subunit distribution in control and mdx4^cv^ skeletal muscles.

#### Histological and immunofluorescence based analyses

Skeletal muscle cryosections were prepared exactly as described [12]. Briefly, 10 micron-thick cryosections were cut from muscle mid bellies on a Leica 1850 cryostat. Cryosections were then stained with hematoxylin and eosin using standard methods. Images of whole hematoxylin and eosin stained cryosections were generated using an automated capture and tiling system comprised of a DP80 digital camera mounted on a BX50 upright microscope (Olympus) fitted with a motorized x-y stage (Prior Scientific) run by CellSens software (Olympus).

For antibody-based immunolabeling, cryosections were fixed in 1 % paraformaldehyde, blocked with 3 % bovine serum albumin in PBS, and then sequentially labeled with primary and secondary antibodies. We used primary antibodies against collagen type I (#ab34710), collagen type III (#ab7778), collagen type IV (#6586), fibronectin (#ab2413) purchased from Abcam and CD68 (#MCA1957GA) from Bio-Rad. Alexa 488 or Alexa 568 fluor-conjugated donkey anti-rabbit secondary antibodies (Jackson Immunoresearch) were used to detect primary antibodies. Images of whole immunolabeled muscle sections were also obtained with the automated capture and tiling system described above.

Hematoxylin and eosin stained cryosections were used to determine myofiber central nucleation, size and size variability. The fraction of centrally nucleated fibers was calculated from at least four whole muscle sections cut from different depths as described [21]. Minimal feret’s diameter and size variability coefficient (a measure of myofiber diameter variability) of muscle fiber types was determined exactly as described [32].

### Physiological measurements

#### Radiotelemetry

Systolic blood pressure (SBP) was monitored using radiotelemetry of the left carotid artery of conscious, non-restrained mdx mice. Briefly, mice under anesthesia with 2% isoflurane were implanted with a telemetry probe (TA11PA-C10, Data Sciences International). Mice were then allowed to recover from surgery over 2 weeks. A cross-over study was performed where one cohort of mice was placed on a diet consisting of BAY 41-8543 (150 ppm) and the other on normal chow for one week, and SBP was measured continuously for the last 2-3 days. Following this, the diets were swapped and SBP was measured again.

#### In situ skeletal muscle force measurements

Tibialis anterior muscle contractility was evaluated with an in-situ muscle test system (Aurora Scientific, Inc.) in Avertin anesthetized mice as described [32]. Briefly, anesthetized mice were positioned on a heated stage and had their hind limb pinned at the knee and the distal tendon of the tibialis anterior secured to the lever arm of a servomotor. The tibialis anterior was stimulated through the peroneal nerve by platinum needle electrodes and adjusted to an optimal length (Lo) for maximum tetanic force generation. Specific force was determined by normalizing tetanic force output to muscle cross-sectional area (Lo□×□pennation angle correction factor□×□density]/muscle mass) at stimulation frequencies ranging from 10 to 200 Hz. To test fatigue susceptibility, tibialis anterior muscles were subjected to a 4-minute fatigue protocol involving 120 isometric tetanic stimulations (200□Hz) at 2-s intervals. Following the fatigue protocol, muscle force recovery was evaluated by determining muscle force output at 1, 3, and 5□minutes. To test the impact of sGC stimulation on mdx muscle susceptibility to lengthening contraction-induced injury, tibialis anterior muscles were subjected to 10 lengthening (20 % beyond Lo) maximal tetanic contractions.

#### Ultrasonography

High frequency, high resolution ultrasonography performed on a Vevo® 2100 high resolution ultrasound imaging platform (VisualSonics) was used to non-invasively measure the amplitude of diaphragm displacement in isoflurane anesthetized mice as described [68]. Amplitude is an *in vivo* measure of diaphragm muscle strength and function.

#### Whole body plethysmography

Ventilatory function was monitored in unrestrained conscious mice using Buxco small animal whole-body plethysmography system and FinePointe software (Data Sciences International). Prior to performing the experiment, single mice were placed in chambers for 1 hour to acclimate daily for 3 consecutive days. For experiments, mice were placed in individual chambers and allowed 10 min to acclimate; then respiratory parameters were measured continuously for the next 10 minutes. The software averaged parameter output generating a mean value every minute for 10 minutes. Parameters recorded included: respiratory frequency (breaths/min), body weight normalized tidal (V_T_; mL/g) and minute volume (V_E_; mL/min/g) and breath rejection index.

#### Echocardiography

Cardiac function in aged mice was evaluated by a Vevo2100 imaging system (VisualSonics) with a MS400 linear array transducer as described [69]. Mean parameter values were analyzed with VevoLab 1.7.1 software (VisualSonics). Briefly, mice were anesthetized with 4% isoflurane and maintained with 1% isoflurane. Depilatory cream was used to remove chest fur and ultrasonographic acoustic gel was used to couple the scan head and skin. Following anesthesia, the mice were fixed in a supine position on a pad with an integrated temperature sensor, a heater and ECG electrodes. Heart rate was monitored continuously and body temperature maintained at 37°C. Short-axis M-mode images were obtained by placing an M-mode gate at mid-papillary muscle level. Cardiac parameters measured in M-mode included: interventricular septum (IVS) thickness (d, s), left ventricular anterior wall (LVAW) thickness (d, s), left ventricular internal diameter (LVID, d and s), left ventricular posterior wall (LVPW) thickness (d, s); left ventricle volume (d, s), stroke volume (SV), cardiac output (CO); % ejection fraction (%EF), % fractional shortening (%FS); LV mass; LV mass corrected. Mitral valve (MV) inflow was assessed from the apical 4 chamber view. MV measurements performed were: early wave peak (E), atrial wave peak (A), aortic ejection time (AET), isovolumic relaxation time (IVRT), isovolumic contraction time (IVCT), E wave deceleration time (DT). Myocardial performance index was calculated as (IVRT + IVCT)/AET as described [53]. Tissue Doppler Imaging (TDI) was performed on mitral valve leaflet tips to measure the velocity of myocardial motion. From TDI we measured the E’ wave corresponding to the motion of the mitral annulus during early diastolic filling of the LV, and the A’ wave which originates from atrial systole during late filling of the left ventricle.

#### Statistics

All data are reported as mean with standard deviation with the exception of Figure 6E and Supplemental Fig 2C. Researchers were blinded to treatment with mouse or sample identity masked during experimental analysis. Statistical outliers were identified by the robust nonlinear regression and outlier removal (ROUT) method using GraphPad Prism software. The specific statistical tests used to determine significant differences between group means are included in figure legends and were also performed in GraphPad Prism with p values < 0.05 considered significant.

## Notes

### Competing Interest Statement

The authors have declared no competing interest.

