## Supplementary Figure Legends for "Soluble guanylate cyclase stimulation mitigates skeletal and cardiac muscle dysfunction in a mdx model of Duchenne muscular dystrophy"

**Supplementary Figure 1**. Western blots of sGC subunit expression. Western blots of β1 and α1 sGC subunit expression in gastrocnemius (A and B) and diaphragm (C and D) muscles from 15-week-old wild type, untreated and BAY41-8543 treated mdx mice. Loading controls are total protein visualized on the immunoblot membrane using Bio-Rad stain-free technology. Red color in B represents saturated pixels. (E) Densitometry quantitation of α1 sGC subunit expression in gastrocnemius and diaphragm muscles showed that α1 sGC subunit expression was similar between all groups.

**Supplementary Figure 2**. Impact of BAY41-8543 dosing regimen on food intake, body mass and blood pressure in mdx mice. (A) Steady state cGMP levels in lungs from wild type (used as reference value), untreated mdx and mdx mice treated for three months with control chow or chow compounded with BAY41-8543. (B) Average mass of chow eaten per mouse per day over the treatment period. (C) Growth curves of wild type, untreated and BAY41-8543 treated mdx mice during treatment period. Growth data were fitted with nonlinear second order polynomial curves. R2 curve fit values for wild type, mdx and mdx BAY41-8543 groups are 0.91, 0.92 and 0.90, respectively. Cohort size for wild type, mdx and mdx BAY41-8543 groups is 10, 13 and 14, respectively. (D) Mean body mass ± SEM of wild type, untreated and BAY41-8543 treated mdx mice at the end of the treatment period. (E) Radiotelemetry assessment of the impact of BAY41-8543 on systolic blood pressure (SBP). Mdx mice were fed control chow or BAY41-8543 supplemented chow ad libitum for one week and SBP was measured continuously over the last 2-3 days. (F) Mean SBP in untreated and BAY41-8543-treated mdx mice. SBP was modestly lowered in treated mdx mice compared with mdx controls during the active night period only. For A, B and D, ** p < 0.01, **** p < 0.0001 by one factor ANOVA with Tukey’s multiple comparison test. For C, wild type versus mdx: * p < 0.05, ** p < 0.01 by repeat measures ANOVA with Bonferroni post hoc test. Wild type versus mdx BAY41-8543: # p < 0.05, ## p < 0.01, ### p < 0.001 by repeat measures ANOVA with Bonferroni post hoc test. For F, * p < 0.05 by t-test.

**Supplementary Figure 3**. sGC stimulation reduces fibronectin positive lesions and promotes fibronectin remodeling to a wild type distribution in the mdx diaphragm. (A) Representative wide field immunofluorescence images of whole diaphragm muscle sections from wild type, untreated and BAY41-8543-treated mdx mice immune-labeled with anti-fibronectin antibodies. Fibronectin-positive macrophage infiltrated muscle lesions are identified with yellow hatched boxes. Experimentalists blinded to sample identity successfully identified treated fibronectin labeled mdx diaphragms 87.5 % of the time. Scale bar 500 microns. (B) Representative wide field immunofluorescence images of diaphragm sections immuno-labeled with an anti-collagen III antibody. Scale bar 100 microns. (C) Western blotting analysis of the expression of α-smooth actin protein, a marker of activated myofibroblasts that participate in interstitial fibrosis. * p < 0.05 by one factor ANOVA with Tukey’s multiple test correction.

**Supplementary Figure 4.** sGC stimulation and increased cGMP have no impact on utrophin expression or mitochondrial content in mdx diaphragm muscles. Western immunoblot and densitometry quantitation of the impact of BAY41-8543 treatment on (A) the expression of protein subunits of mitochondrial respiratory chain complexes I-V and (B) mitochondrial marker VDAC1. (C) Western immunoblot and densitometry quantitation of the impact of BAY41-8543 treatment on utrophin expression in mdx diaphragm muscle. *** p < 0.001 and **** p < 0.0001 by one factor ANOVA with Tukey’s multiple test correction.

**Supplementary Figure 5**. sGC stimulation has no impact on the in situ isometric twitch properties of tibialis anterior muscles from mdx mice. As expected, peak twitch force was increased in mdx mice relative to wild type controls. BAY41-8543 had no effect on peak twitch force in mdx mice. (B) As expected, specific twitch force was reduced in mdx mice compared to wild type mice. BAY41-8543 had no effect on specific twitch force in mdx muscle. (C) The time taken to half relaxation (50 % of peak force) was similar between wild type, untreated and BAY41-8543-treated mdx mice. (D) The time taken for the generation of peak twitch force was similar between wild type, untreated and BAY41-8543 treated mdx mice. *** p < 0.001, **** p < 0.0001 by one factor ANOVA with Tukey’s multiple comparison correction.

**Supplementary Figure 6.** sGC stimulation in wild type mice mimics altered respiratory function in mdx mice. (A-K) Whole body plethysmography analysis of breathing function in 19-20-week-old wild type and mdx mice and mdx mice treated since 3-4 weeks of age with BAY41-8543. (A) Inspiratory time T_I_. (B) Expiratory time T_E_. (C) Breathing cycle duration, which is the sum of inspiratory and expiratory times. (D) Estimated peak inspiratory flow (PIF). (E) Estimated peak expiratory flow. (F) Expiratory airflow rate when 50 % of the air volume has been exhaled during normal breathing (EF50). (G) Relaxation time-the time is takes to expire a certain percentage (65 %) of tidal volume. (H) Duration of breaking or the percentage of the breath occupied by the transition from inspiration to expiration. (I) Duration of pause or the percentage of the breath occupied by the transition from expiration to inspiration. (J) Pau (pause) is the ratio PEF/PIF and is an indication of bronchoconstriction. * p < 0.05, ** p < 0.01, *** p < 0.001, **** p < 0.0001 by two factor ANOVA with Tukey’s multiple comparison correction.

**Supplementary Figure 7**. sGC stimulation impact on cardiac geometry and dysfunction in aged mdx mice assessed by echocardiography. (A) Percent fractional shortening. (B) Left ventricular anterior wall (systole). (C) Left ventricular anterior wall (diastole). (D) Left ventricular posterior wall (systole). (E) Left ventricular posterior wall (diastole). (F) Left ventricular internal diameter (systole) (G) Left ventricular internal diameter (diastole). (H) Isovolumic relaxation time shown here again for reference (see Figure 8K). (I) isovolumic contraction time. (J) Aortic ejection time. (K) Myocardial performance index. Group sizes for wild type, mdx and BAY41-8543-treated mdx are 5-7, 8 and 6, respectively. Unpaired t-test where # and ## represents p < 0.05 and p < 0.01 for wild type versus mdx, respectively. * and ** represents p < 0.05 and p < 0.01 for mdx versus mdx BAY41-8543, respectively. φ represents p < 0.05 for wild type versus mdx BAY41-8543.
