## Supplementary figures and images for "Soluble guanylate cyclase stimulation mitigates skeletal and cardiac muscle dysfunction in a mdx model of Duchenne muscular dystrophy"

### Supplementary Figure 1

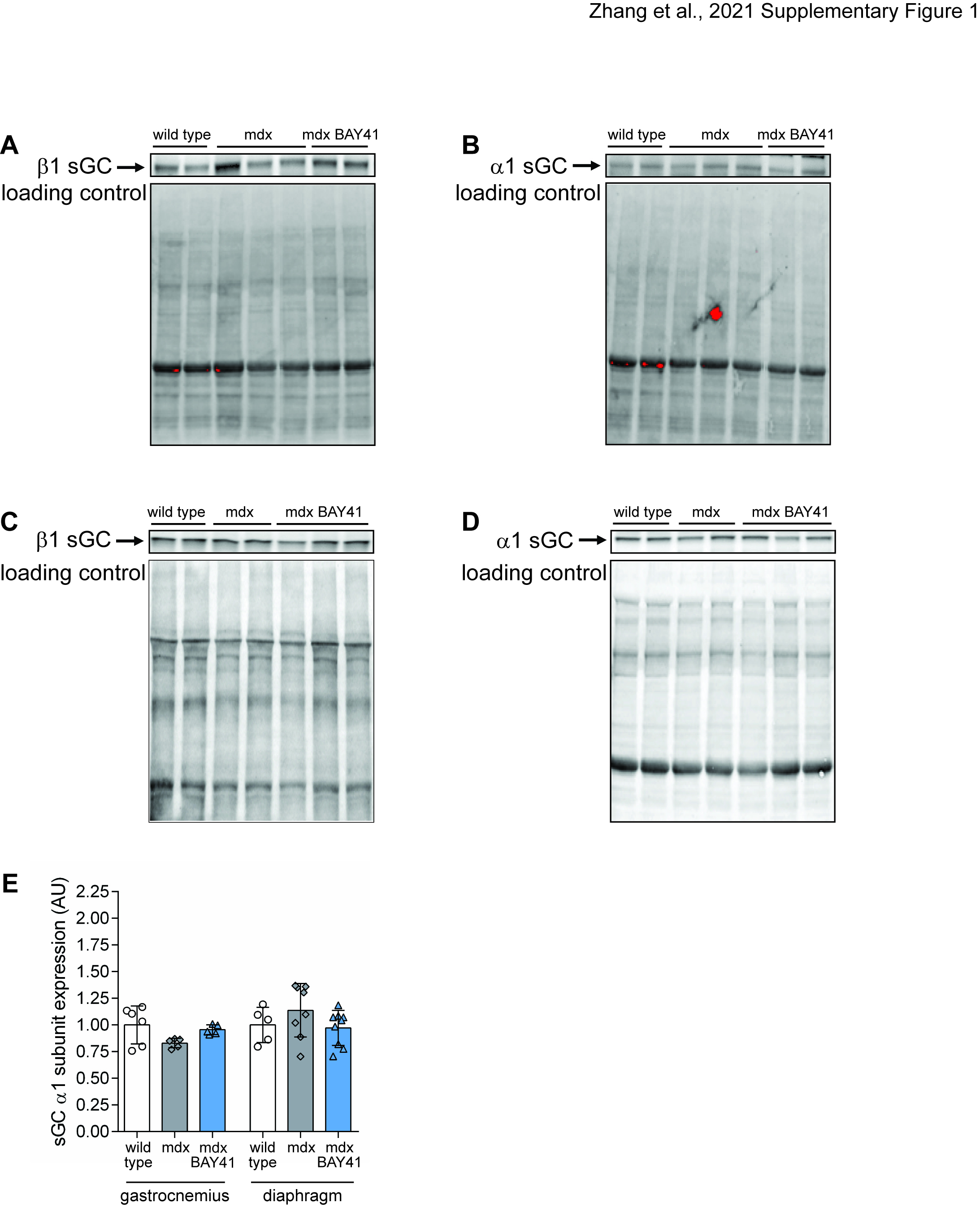

### Supplementary Figure 2

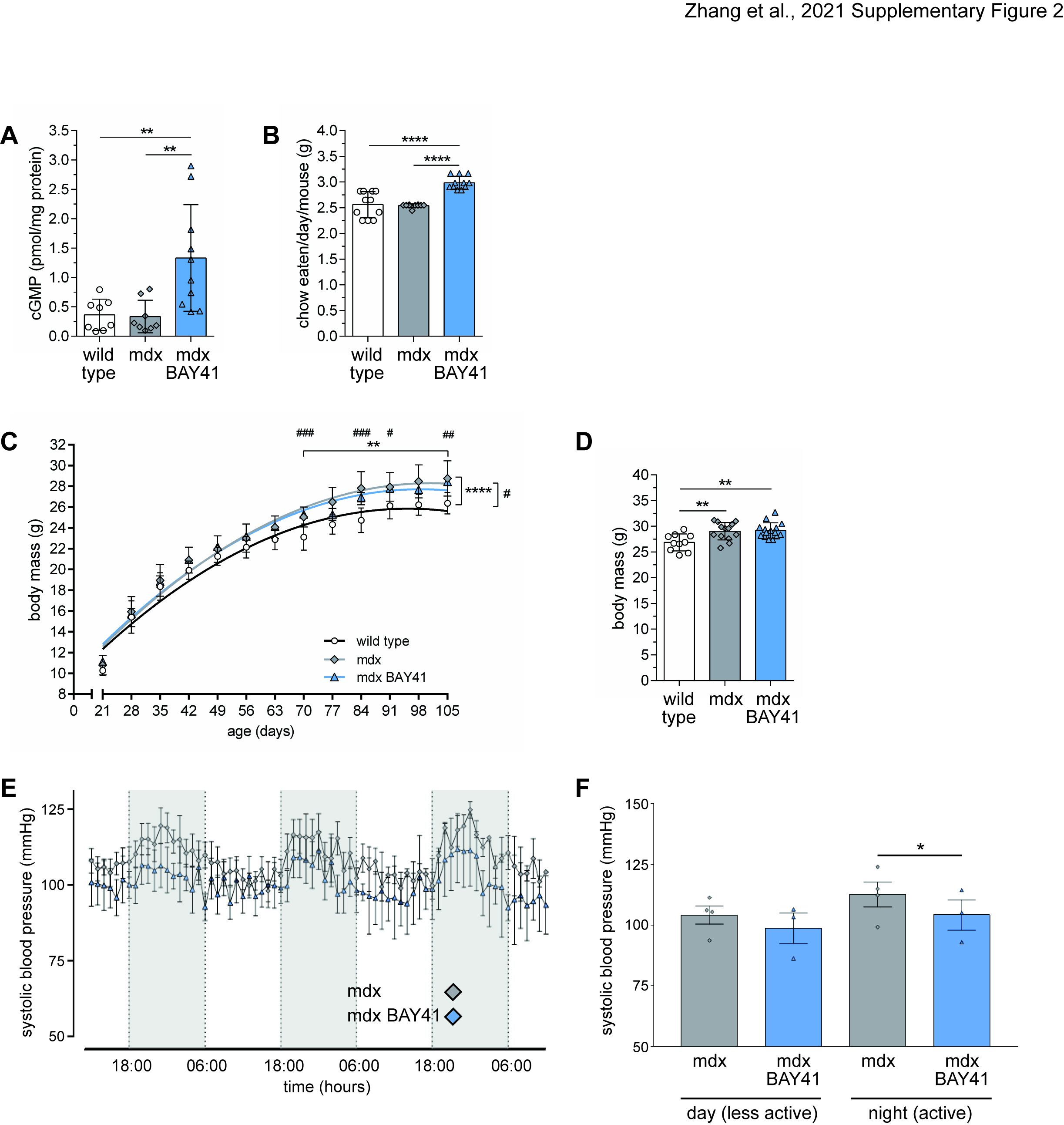

### Supplementary Figure 3

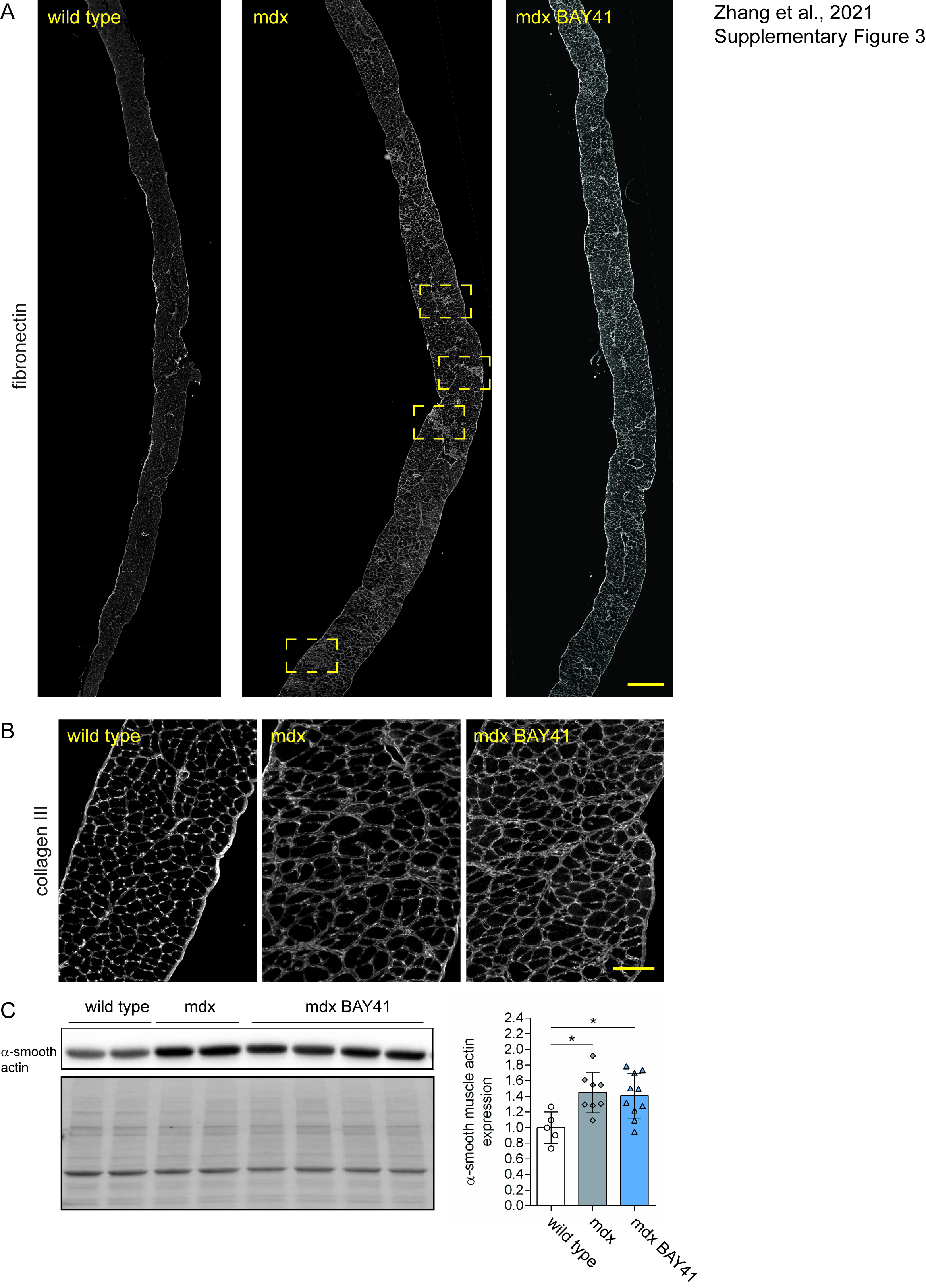

### Supplementary Figure 4

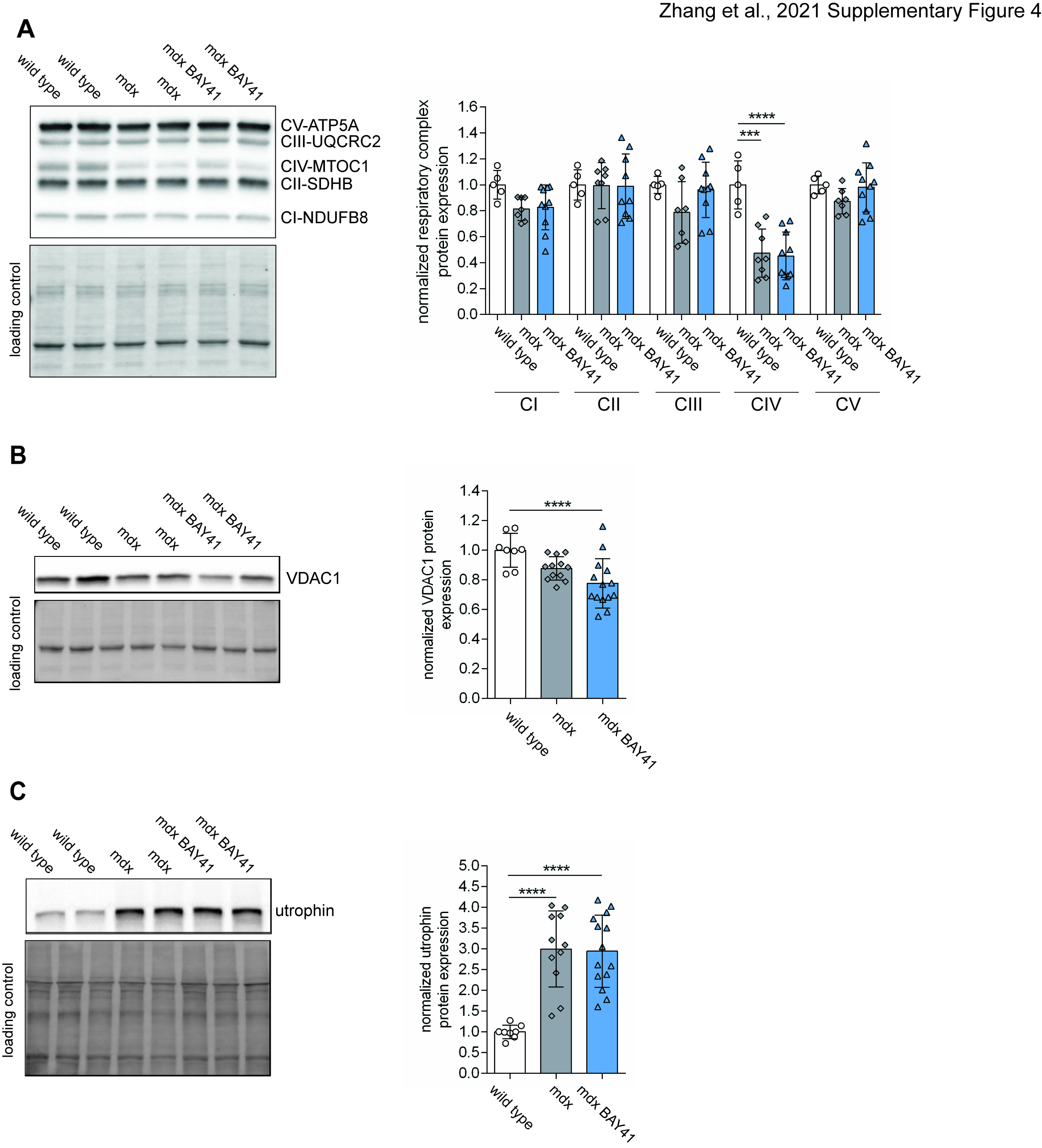

### Supplementary Figure 5

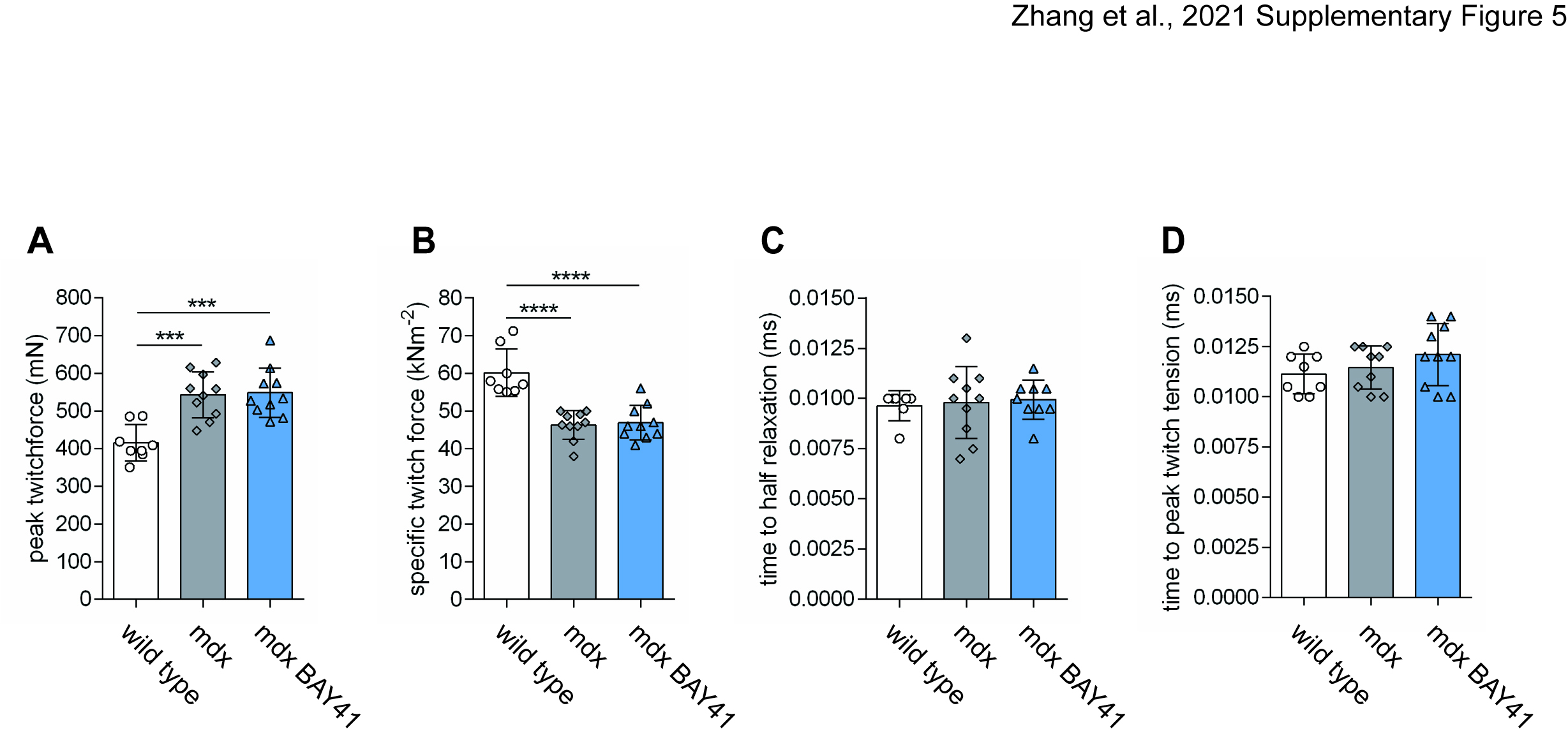

### Supplementary Figure 6

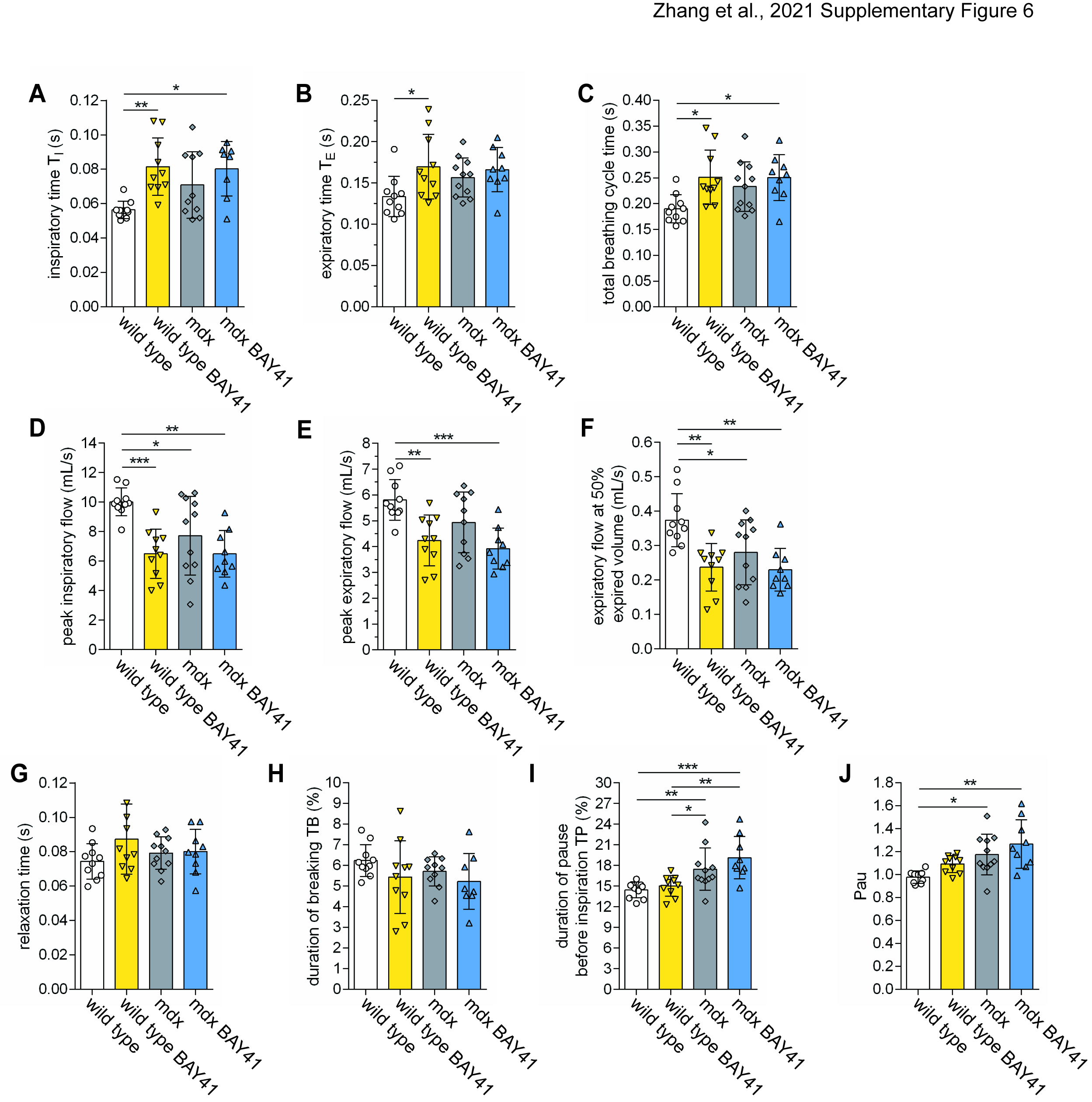

### Supplementary Figure 7

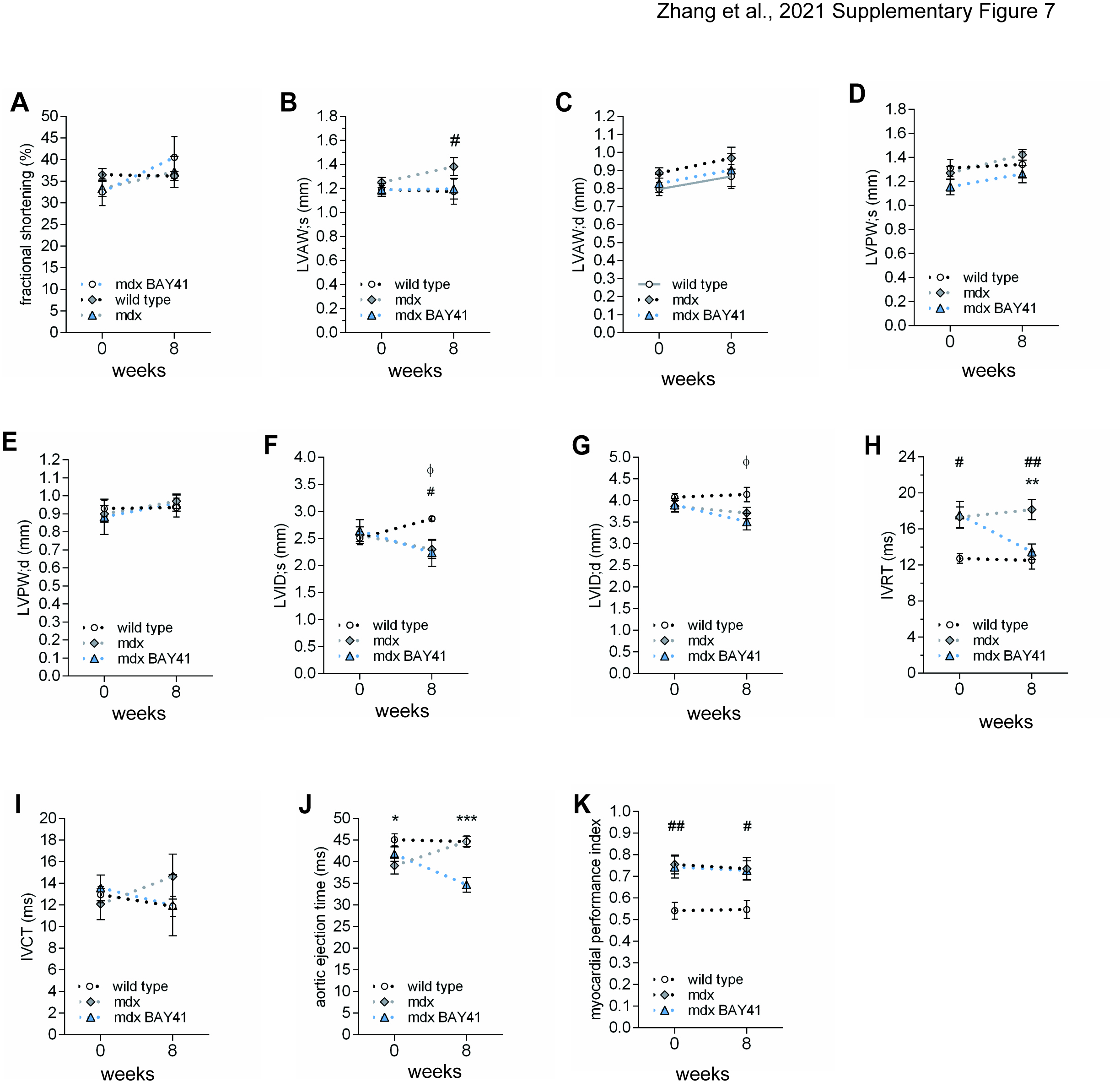
